# An Eutherian-Specific microRNA Controls the Translation of *Satb2* in a Model of Cortical Differentiation

**DOI:** 10.1101/2020.10.26.355214

**Authors:** Manuella Martins, Silvia Galfrè, Marco Terrigno, Luca Pandolfini, Irene Appolloni, Keagan Dunville, Andrea Marranci, Milena Rizzo, Alberto Mercatanti, Laura Poliseno, Francesco Morandin, Marco Pietrosanto, Manuela Helmer-Citterich, Paolo Malatesta, Robert Vignali, Federico Cremisi

## Abstract

Cerebral cortical development is controlled by key transcription factors that specify the neuronal identities in the different cortical layers. These transcription factors are crucial for the identity of the different neurons, but the mechanisms controlling their expression in distinct cells are only partially known. Here we investigate the expression and stability of the mRNAs of Tbr1, Bcl11b, Fezf2, Satb2 and Cux1 in single developing mouse cortical cells. We focus on Satb2 and find that its mRNA expression occurs much earlier than its protein synthesis and in a set of cells broader than expected, suggesting an initially tight control of its translation, which is subsequently de-repressed at late developmental stages. Mechanistically, *Satb2* 3’UTR modulates protein translation of GFP reporters during mouse corticogenesis. By *in vitro* pull-down of *Satb2* 3’UTR-associated miRNAs, we select putative miRNAs responsible for SATB2 inhibition, focusing on those strongly expressed in early progenitor cells and reduced in late cells. miR-541, an Eutherian-specific miRNA, and miR-92a/b are the best candidates and their inactivation triggers robust and premature SATB2 translation in both mouse and human cortical cells. Our findings indicate that RNA interference plays a major role in the timing of cortical cell identity and may be part of the toolkit involved in specifying supra-granular projection neurons.

## INTRODUCTION

The mammalian neocortex consists of six cell layers (I-VI) generated by radial migration of neuroblasts following an inside-out mechanism (Greig et al., 2013). Glutamatergic projection neurons are formed after the generation of layer I neurons in two main neurogenetic waves: deep projection neurons (DPNs) of layers V-VI are generated first, followed by superficial projection neurons (SPNs) of the supragranular layers II-III (Figure 1A). Generation of layer IV neurons follows the generation of DPNs and precedes SPNs formation. Proper regulation of this developmental process is crucial and its impairment results in various disorders such as brain malformations or psychiatric diseases (Sun and Hevner, 2014). The capability to generate distinct classes of neurons depends on the progenitor cell cycle state and neuron birth date (McConnell and Kaznowski, 1991). Epigenetic birthmarks may regulate the ability of cortical progenitor cells to establish neuron identity already in the first hour following the last cell division (Telley et al., 2019). After this, the expression of a few cell identity transcription factors (CITFs) is necessary to impart distinct cell fates, with TBR1, BCL11b, FEZF2, SATB2 and CUX1 playing an important role among them (Alcamo et al., 2008; Cubelos et al., 2010; Hevner et al., 2001, 2003; Leone et al., 2015; Srinivasan et al., 2012). These factors may initially establish early mutual activating or repressive interactions; beyond this early phase, depending on the cell context and the timing of corticogenesis, some of these interactions may change and combinatorial action may ensue to refine terminal cell phenotype (Alcamo et al., 2008; Britanova et al., 2008; Chen et al., 2008; Harb et al., 2016; Jaitner et al., 2016; McKenna et al., 2015). A precise timing of expression of these and other factors is required to ensure appropriate differentiation of the neocortex. The exact mechanisms dictating the timely expression of CITFs in one given progenitor cell and its progeny is still under scrutiny.

**Figure 1.**
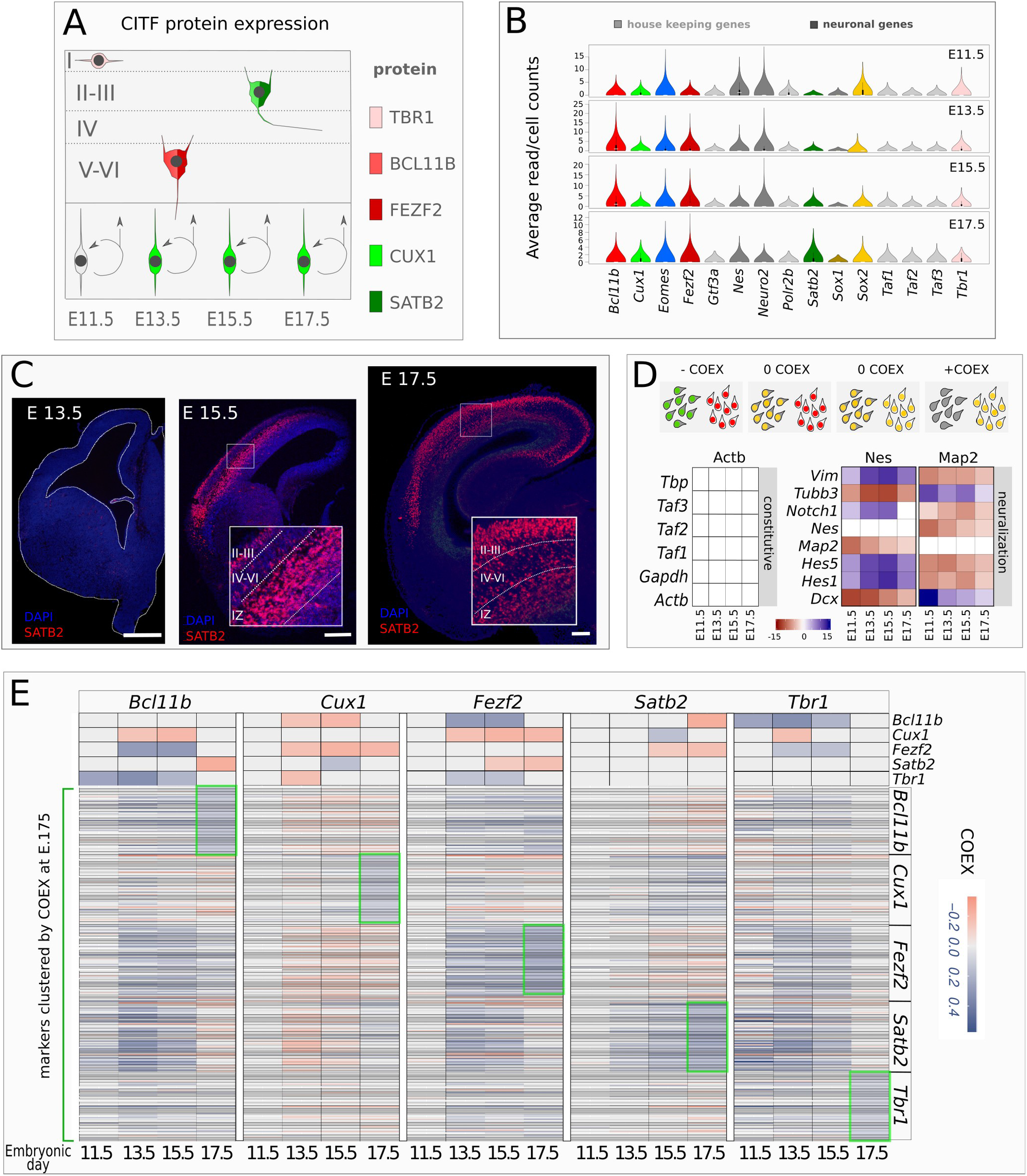
CITF expression analysis. A, Simplified outline of cortical layering. E11.5-E17.5, embryonic day 11.5-17.5. Layers are labeled by Roman numerals. B, Violin plots show average raw counts/cell of genes indicated in labels. Constitutive gene are in light grey. C, Coronal sections of mouse embryonic brain showing SATB2 immunodetection at different embryonic (E) developmental times of corticogenesis. Roman numerals indicate cortical layers. IZ, intermediate zone. D, Top schematic shows COTAN COEX relation to the pattern of expression of two genes (red and green) in single cell. Bottom shows COEX values of pairs of constitutive genes (left matrix) or neural differentiation markers (right matrix) at the different developmental times shown in labels. E, COTAN COEX values of CITFs and genes associated to them by high COEX at E17.5. The top side of the matrix shows the COTAN COEX relation between pairs of CITFs. The bottom part of the matrix reports COTAN COEX values between distinct CITFs and the genes that are more highly co-expressed with each of them at E 17.5 (green boxes).

The evolution of the mammalian cortex is characterized by the progressive thickening of the supragranular cell layer(s) (Dehay and Kennedy, 2007; Dehay et al., 2015). A sudden evolutionary change during mammalian cortex evolution may be the heterochronic appearance of the cortical transcription factor SATB2 with respect to the corresponding mRNA. Indeed, it was recently shown that SATB2 protein expression is delayed in Eutherians compared to Metatherians and such delay seems responsible for the development of the inter-hemispheric callosal connections generated from the supra-granular cells in Eutherians (Paolino et al., 2020). After its evolutionary appearance, the continuous expansion of the corpus callosum (CC), and of the supra-granular cell layer it stems from, represents the distinguishing feature of the placental neocortex, including that of higher primates. Notably, in higher primates SATB2 protein appearance is delayed over an extended period, possibly crucial for supra-granular cell layer expansion (Otani et al., 2016). In this aspect, the control of developmental timing of SATB2 during cortical neurogenesis may be of crucial importance. In this paper, we have first investigated the differential stability of mRNAs for key CITFs involved in mammalian corticogenesis, namely Bcl11b, Cux1, Tbr1, Fezf2 and Satb2, by exon/intron stability analysis (EISA) (Gaidatzis et al., 2015). We find that among them only Satb2 mRNA shows an increase in exon/intron (E/I) ratio due to an improved stability and rate of its transcription. We then show that a post-transcriptional control is played by microRNAs (miRNAs) acting on Satb2 3’UTR. We isolated miRNAs that bind to this region and focus on miR-541, a new, Eutherian specific miRNA; we show that miR-541 delays, both in vivo and in vitro, SATB2 protein production with respect to Satb2 mRNA transcription. We discuss the potential implications of miR-541 action in the scenario of cortical evolution.

## RESULTS

### Satb2 is co-transcribed with other CITFs in early cortical cells before its translation

Since DPNs and SPNs are sequentially generated in an inside-out fashion from embryonic day 11.5 (E11.5) to E17.5 in mouse (Figure 1A), we expect that the mRNA of CITFs is regulated in selected progenitor cells during this time window and tested this assumption by re-analyzing single-cell RNA sequencing (scRNA-seq) datasets of mouse cortex at E11.5, E13.5, E15.5, E17.5 (Yuzwa et al., 2017). We compared the average expression level of the 5 CITFs, evaluated as raw counts/cell, to the that of transcription factors with constitutive expression (Figure 1B). The mRNA expression levels of all 5 CITFs are comparable to those of constitutive transcription factor genes since E11.5, indicating that the expression of these 5 mRNAs could have a biological relevance already at very early stages of corticogenesis. However, we did not detect SATB2 translation at E 13.5 (Figure 1C). Although a minority of SATB2-positive cells were reported at 13.5 (Alcamo et al., 2008; Britanova et al., 2008), reliable onset of SATB2 protein expression was not described earlier than E14 (Paolino et al., 2020), suggesting a post-transcriptional regulation of Satb2 mRNA. To get insight on the mechanism of CITF transcriptional activation in specific cell subsets we analyzed CITF co-expression in single cells by Co-expression Table Analysis (COTAN) (Galfrè et al., 2020; Galfrè and Morandin, 2020).

COTAN can assess the co-expression of gene pairs in a cell and, by extending this analysis to all gene pairs in the whole transcriptome, it can infer the tendency of a gene to be constitutively expressed, or expressed in a subset of differentiating/differentiated cells. Positive coexpression index (COEX) denotes the co-expression of two genes, while negative COEX indicates disjoint expression; COEX near 0 is expected if one or both are constitutive genes (Figure 1D, top) or when the statistical power is too low. Accordingly, our analysis gives COEX values close to zero for constitutive mRNA pairs (Figure 1D, left; Supplementary files 1-8). Conversely, high co-expression (positive COEX) is found for mRNA pairs of known molecular markers of neural progenitor cells (*Nestin, Vimentin*, *Notch1, Hes1-5)* or postmitotic cells and differentiating neurons (*Dcx, Tubb3, Map2*). Finally, disjoint expression (negative COEX) is detected between mRNA pairs of these two groups at all developmental stages (Fig. 1D). All CITFs show reciprocal mRNA co-expression patterns consistent with their known protein expression pattern in different cell types, except *Satb2,* whose COEX with each of the other 4 CITFs is comparable to that of constitutive genes at E11.5 and E13.5 (compare Figure 1A and E, top).

We considered the genes most highly co-expressed with each CITF gene (Figure 1E bottom, Supplementary Figure 1) at E17.5. At this stage, the final pattern of co-expression of each CITF gene with co-clustered markers (green boxes in Figure 1E) differs from the patterns at earlier stages of corticogenesis (Figure 1E). This suggests that initially CITF gene expression is not cell layer-specific, but that cell-specific CITF gene expression is reached towards the end of layer formation.

COTAN Gene Differentiation Index (GDI) discriminates between constitutive and non-constitutive genes by globally integrating COEX values (Galfrè et al., 2020) (Figure 2A). We used GDI analysis to infer the propensity of CITFs to be expressed in restricted cell subsets during corticogenesis. Notably, the global relation between GDI and mRNA level values (Figure 2B), and the global GDI distribution (Figure 2C), are comparable during the four developmental times included in the analysis.

**Figure 2:**
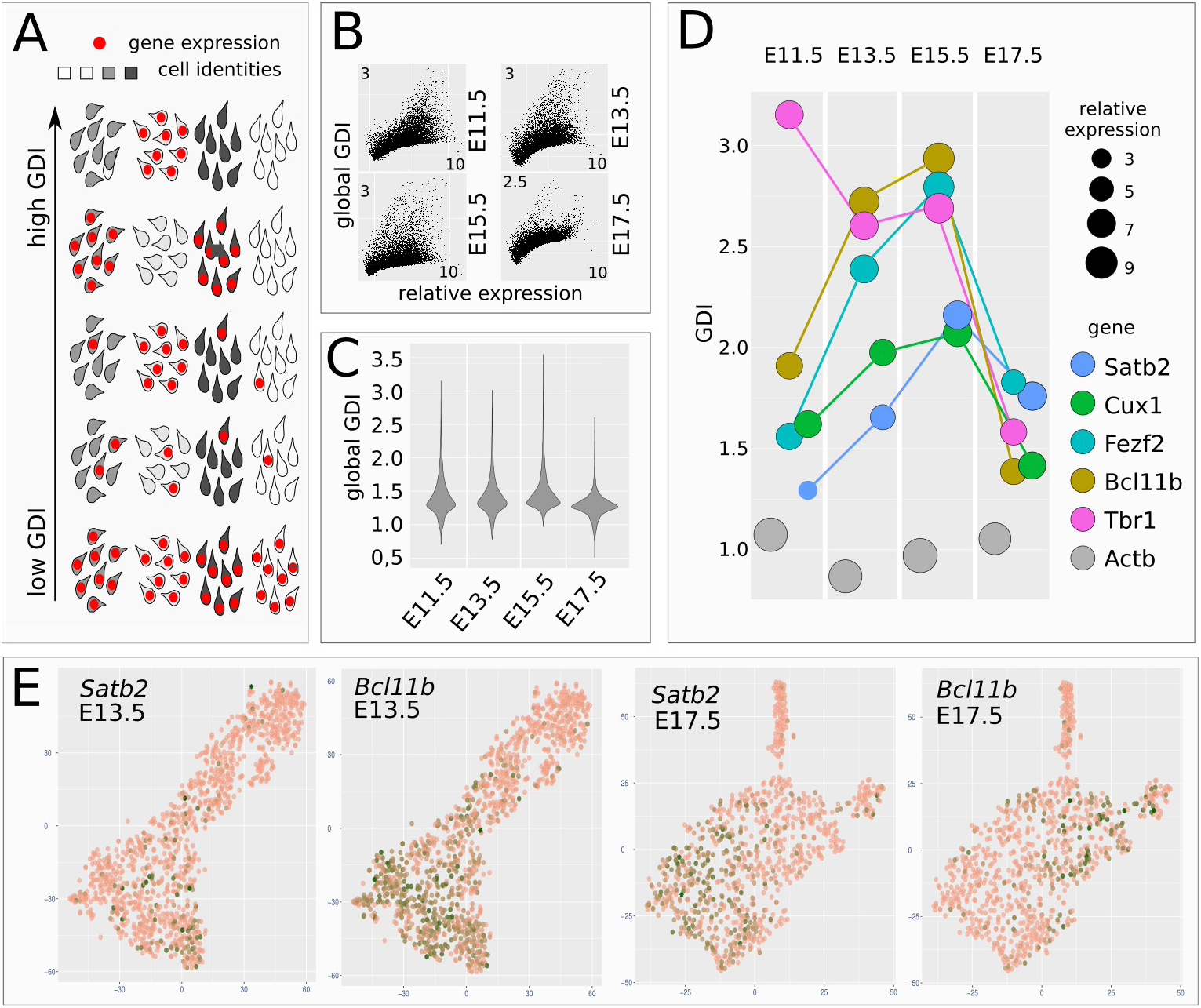
Figure 2 CITF transcription in distinct cell clusters. A, Schematic shows how GDI can infer the degree of gene pair co-expression in cell populations with different cell identities. B, plots show GDI and gene mRNA expression levels at different developmental times. C, violin plots report global GDI distribution during corticogenesis. D, Distinct CITFs show different GDI according to their translational onset. E, t-SNE clustering of early (DIV13.5), or late (DIV17.5) cells. Panels show read distribution of the indicated gene on cell clusters.

This observation supports the use of GDI analysis to evaluate whether a mRNA species changes its pattern of cell distribution during corticogenesis, and becomes restricted to a particular cell lineage/ layer. Unlike constitutive genes as *Actb*, CITFs showed marked GDI changes during corticogenesis (Figure 2D). *Tbr1* mRNA shows a peak at E11.5, consistent with early localized TBR1 protein expression in layer 1 neurons (Hevner et al., 2001). *Bcl11b* and *Fezf2*, followed by *Satb2* and *Cux1*, increase their GDI until E15.5 paralleling their respective onset of protein expression (compare Figure 2D with Figure 1A).

The drop of GDI observed at E17.5 correlates with, and might be explained by, the increased heterogeneity of the cell types co-expressing different combinations of CITF proteins at the end of corticogenesis (Lodato and Arlotta, 2015), although it may also be due to post-transcriptional CITF regulation. Notably, *Satb2* displays the lowest GDI levels among CITFs at E11.5-13.5, when its protein is not yet detectable, suggesting that post-transcriptional mechanisms account, at least in part, for the subsequent restricted expression of SATB2 protein in SPNs.

Finally, we used a conventional t-SNE analysis of gene expression on cells clusters (Figure 2E). The lack of a cell-type restricted distribution of *Satb2* mRNA at early stages is also suggested by its partial overlap with *Bcl11b* mRNA in E13.5 cell clusters, as compared to E17.5 clusters.

### *Satb2* 3’UTR drives RISC-dependent translational inhibition in early cortical cells

We then took advantage of Exon-Intron Split Analysis (EISA) (Gaidatzis et al., 2015; La Manno et al., 2018) to verify whether a time-dependent instability of *Satb2* mRNA could account for the inability to detect SATB2 protein at E13.5, when *Satb2* transcription is already robust and apparently spread in different cell clusters. EISA can evaluate if a mRNA species changes its stability during developmental processes, assuming that the intronic sequences are rapidly spliced and that their levels reflect the gene transcriptional rate (see schematic in Figure 3A, left). Because layer identity commitment is established before neuron birth date (McConnell and Kaznowski, 1991; Telley et al., 2019), we analyzed RNA-seq datasets of progenitor cells (Chui et al., 2020). We observed that *Satb2* Exon/Intron (E/I) ratio significantly increases from E11.5 to E17.5, *Bcl11b* E/I increases from E11.5 to E13.5 and *Fezf2* E/I increases from E13.5 to E15.5, while the E/I of the other CITFs and of *Actb* shows no significant changes (Figure 3A, middle panel). Notably, *Satb2* E/I ratio increase is paralleled by a dramatic increase of its transcription levels from E11.5 to E17.5 (Figure 3A, right), as measured by intron read abundance, making its E/I increase more relevant than that of *Bcl11b* and *Fezf2*. *Satb2* E/I fold change between E13.5 and E15.5 settles in the highest quartile of the E/I increase (Figure 3B, Supplementary file 9), suggesting high biological relevance and supporting close relationship between the increase of *Satb2* mRNA stability and the onset of SATB2 translation. We thus focused our attention on the post-transcriptional regulation of *Satb2*.

**Figure 3.**
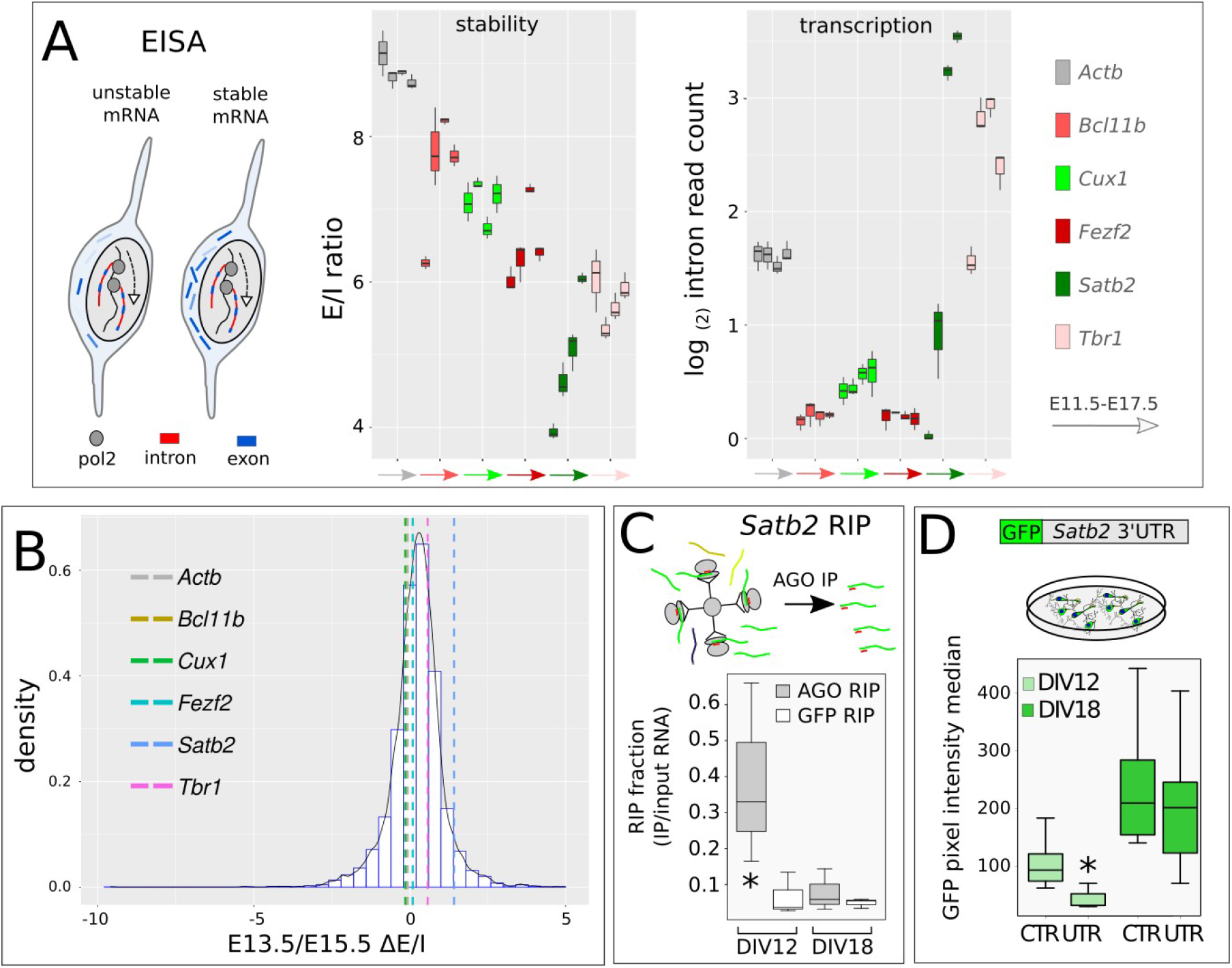
Cortical mRNA exon/intron analysis and SATB2 translational inhibition. A, Exon-Intron split analysis (EISA) of CITF mRNAs. Outline shows different ratios of exonic and intronic sequences in relation to mRNA stability as rationale at the basis of EISA. Box plots show the ratio of exon/intron (E/I) read counts, and intron read counts, for distinct CITFs and *Actb* (constitutive control gene) in cortical progenitors at different *in vivo* embryonic times. B, Density plot of Exon/Intron (E/I) ratio fold change between E13.5 and E15.5. C, qRT-PCR evaluation of Argonaute (AGO)-interacting *Satb2* mRNA. Values on Y axis report the ratio of RT-PCR-detected, immunoprecipitated *Satb2* mRNA with respect to the input (AGO RIP). GFP RIP, control immunoprecipitation with anti-GFP Ab. N= 3 independent experiments. D, Expression of *Satb2* 3’ UTR-bearing GFP reporter after lipofection in corticalized mESCs. N= 3 independent experiments. Cells were transfected 48 hours before the time of analysis indicated in labels.

We reasoned that changes in *Satb2* mRNA stability could be induced by miRNA-dependent RNA interference. Indeed, by high-throughput analysis of miRNA-mRNA interactions at single cell level, distinct miRNAs were recently associated to functional modules involved in the control of different cortical cell identities (Nowakowski et al., 2018). To gain insights on RNA interference during early corticogenesis, we employed mESCs, whose *in vitro* neural differentiation can be steered to closely reproduce the early stages of cortical development, including time-regulated expression of TBR1, BCL11B and SATB2 protein (Bertacchi et al., 2015; Gaspard et al., 2008). In this experimental system, we measured the enrichment of *Satb2* mRNA after AGO2 immunoprecipitation. By qRT-PCR, a significant enrichment of AGO2-bound *Satb2* mRNA is detected in cells after 12 days *in vitro* (DIV) compared to GFP immunoprecipitation used as control, indicating a relevant miRNA silencing activity at an early stage of *in vitro* corticogenesis (Figure 3C). Notably, we found no enrichment at DIV18, consistent with a significant increase of SATB2-positive cells at this time (Bertacchi et al., 2015).

The dynamic binding capacity of *Satb2* mRNA to AGO2 at different times of development is in line with the ability of *Satb2* 3’UTR to inhibit protein translation in early, but not late, cortical cells. Indeed, at DIV12 the transfection of a GFP reporter carrying *Satb2* 3’UTR yields decreased fluorescence levels compared to control, while at DIV18 the reporter activity is not significantly affected (Figure 3D), consistent with robust SATB2 translation at this late stage (Bertacchi et al., 2015). *Satb2* 3’UTR is able to control translation also *in vivo*, as shown by *in utero* electroporation (IUE) of a GFP reporter/sponge. At stage E13.5, the proportion of SATB2-GFP double-positive cells with respect to GFP-positive cells is significantly higher in a cortex electroporated with a 3’UTR-bearing sensor compared to a control-electroporated cortex (Supplementary Figure 2). These results indicate that *Satb2* 3’UTR can inhibit the translation of its mRNA in early-generated neurons.

### miRNAome time trajectories describe cortical development progression

We then set out to identify miRNA candidates regulating *Satb2* expression. To this aim, we sorted *Sox1*::GFP corticalized mES cells, which are enriched in progenitors, and first analyzed their global miRNA profiles in comparison with the profiles of non neuralized mES cells, post-mitotic corticalized mES cells obtained by AraC treatment, or mouse cortex, at different developmental times (Figure 4A-D, Supplementary file 10). MiRNAome PCA shows high consistency between miRNA profile and cell identity. MiRNAomes of non neuralized mES cells are well separated from those of corticalized mES cells and of cortex, which instead cluster together, confirming that our protocol mimics a genuine cortical identity *in vitro* (Figure 4A). The time of *in vitro* differentiation distributes both neuron (Figure 4B) and progenitor (Figure 4C) miRNAomes along PC3, in agreement with the relative position of E12 and P0 cortex miRNAomes, indicating high conservation of the mechanisms accounting for the timing of layer formation in our *in vitro* conditions. Finally, PC3 discriminates between progenitor and neuron miRNAomes (Figure 4D), indicating that these distinct cell states are maintained throughout the differentiation process.

**Figure 4.**
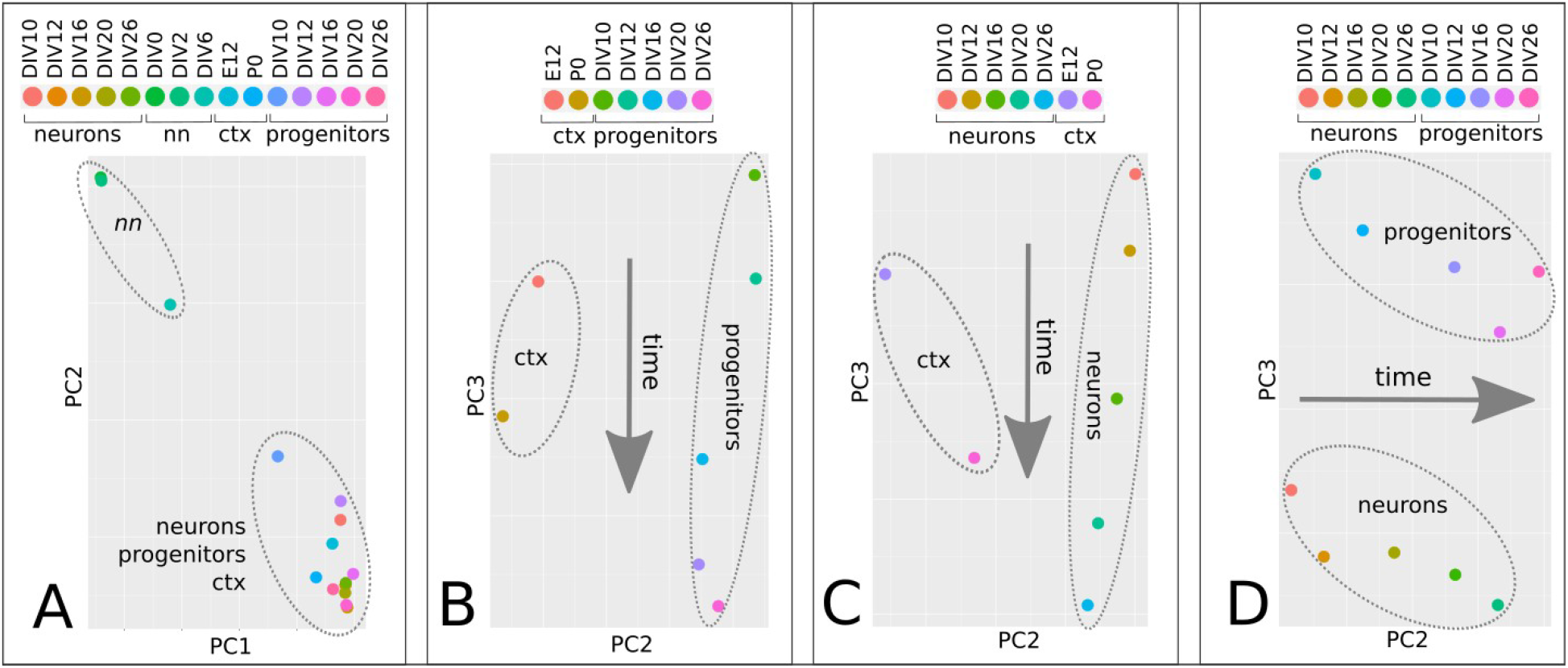
miRNAome time trajectories in corticogenesis. A-C, PCA of miRNA global profiles of non neuralized mES cells (nn), neural progenitor cells (Sox1::GFP corticalized mESCs), post-mitotic cells (Ara-C-treated corticalized mESCs) and mouse cortex (ctx) at different developmental times. Four different combinations of the four above mentioned groups are shown.

### Selected miRNAs directly bind *Satb2* 3’UTR in early cortical cells

To select miRNAs that directly interact with *Satb2* 3’ UTR at DIV12 and DIV18 we employed miR-catch analysis, which is based on the recovery of mRNA/RISC/miRNA complex by digoxigenin-labeled probes complementary to the target mRNA (Marranci et al., 2019; Vencken et al., 2015). This method quantifies bound miRNAs through small RNA-sequencing, measuring miRNA enrichment with respect to the input (total miRNAs) (Figure 5A). With this approach, we found twelve miRNAs that bind to *Satb2* mRNA and are significantly enriched at DIV12; among these, miR-541 and miR-3099 are not enriched at DIV18, thus representing candidates for SATB2 inhibition in early, but not late, cortical cells (Figure 5B, Supplementary Figure 3,4, Supplementary File 11).

**Figure 5.**
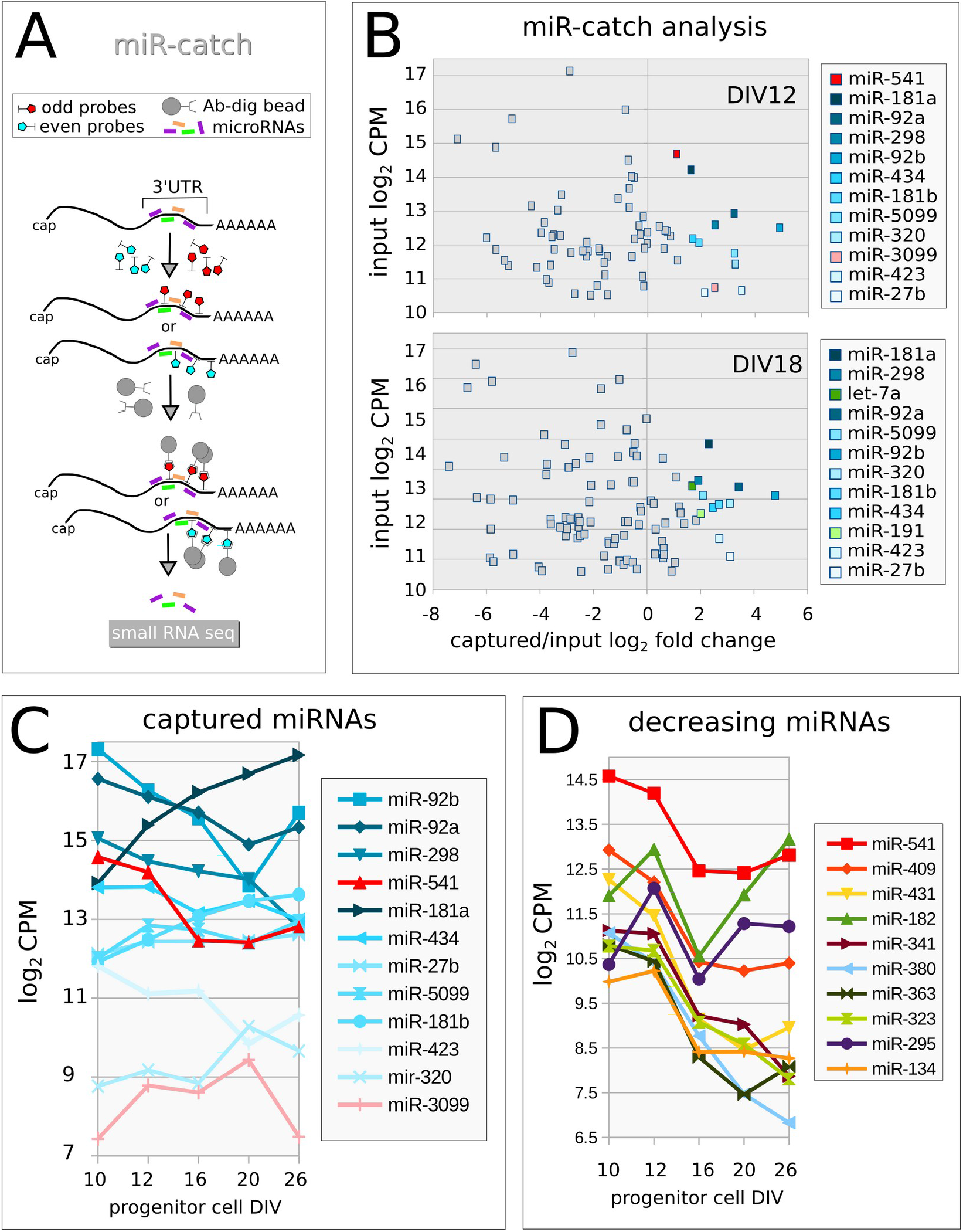
*Satb2* interacting miRNAs. A, Outline of miR-catch method. B, enrichment of captured miRNAs (x-axis) with respect to input (Y axis) at the indicated time. CPM, counts per million. Color labels indicate significantly enriched miRNAs (non-parametric noiseqbio test probability > 0.9) (Tarazona et al., 2015). C, Developmental expression patterns of *Satb2*-captured miRNAs in *Sox1*::GFP progenitor cells. D, developmental expression of miRNAs with the highest monotonic developmental decrease in *Sox1*::GFP progenitor cells. CPM, counts per million.

Because of its extremely low expression levels (Figure 5C) we did not further investigate miR-3099 and focused our attention on the other miRNAs.

We analysed the abundance of the twelve captured miRNAs in progenitor cells and found that only miR-92a-b and miR-541 show robust decrease between DIV12 and DIV16, when SATB2 translation is de-inhibited (Figure 5C). We thus focused our attention on these three miRNAs. miR-92 was already shown to play a major role in inhibiting EOMES (TBR2) translation and preventing early generation of basal progenitors, which give rise to supragranular neurons in mouse (Bian et al., 2013; Nowakowski et al., 2013). Conversely, miR-541 has never been found involved in cortical development. miR-541 belongs to an evolutionary new miRNA cluster (mir-379-mir-410 in mouse, mir-379-656 in humans), which is located into a large miRNA-containing gene (*Mirg*) inside the *DLK-DIO3* locus (Edwards et al., 2008; Glazov et al., 2008; Winter, 2015). *Mirg* orthologues have been found in all Eutherian, which hold inter-hemispheric cortical connections forming the corpus callosum, but not in Metatherian, Prototherian, or any other vertebrates, which lack corpus callosum. Moreover, gene targets for mir-379/mir-656 cluster are significantly over-represented in Gene Ontology terms associated with neurogenesis and embryonic development, and miRNA expression was detected in brain and placenta, suggesting that Mirg appearance was one of the factors that drove the evolution of the placental mammals (Glazov et al., 2008). miR-541 shows an *in vitro* pattern of expression that closely matches the time-dependent inhibition of SATB2 translation and follows a sudden down-regulation between DIV12 and DIV16 (Figure 5 C). In addition, at E13.5 miR-541 is widely expressed in the ventricular zone (VZ), subventricular zone (SVZ) and mantle zone (MZ), when SATB2 protein is undetectable, while at E15.5 the miRNA is expressed in the cortical plate (CP), when the protein is detected in VZ, SVZ, intermediate zone (IZ) and migrating cells (Supplementary Figure 5). Finally, miR-541 developmental decrease is comparable to that of the most heavily downregulated miRNAs from DIV10 to DIV12 (Figure 5D), supporting its candidacy for the control of SATB2 inhibition during the early corticogenesis.

### miR-541 and miR-92a/b inhibit SATB2 translation in both mouse and human early cortical cells

We then inhibited miR-541 and mir_92a/b by transfection of a complementary locked-RNA (antagomiR) in mouse ES corticalized cultures (Figure 6A). This results in a premature and massive increase of SATB2-positive cells compared to control-transfected cells (Figure 6B top, C). Notably, miR-541 has no predicted binding site on *Eomes* 3’UTR; thus, its effect on SATB2 translation is unlikely to depend on increased EOMES translation and consequent induction of basal progenitor identity (Sessa et al., 2008), which may be the case with miR-92a/b inhibition. miRNA inhibition both anticipates the onset of SATB2 protein detection, as indicated by the effect of transfection at DIV10, and increases the efficiency of translation at later time points, as emphasized by the outcome of DIV12 transfection. Finally, we observed a similar effect when downregulating miR-541 and miR-92 in corticalized human induced pluripotent cells (hiPSCs) (Figure 6B, bottom, C), suggesting evolutionary conservation of the mechanism of cell-type-specific SATB2 expression.

**Figure 6.**
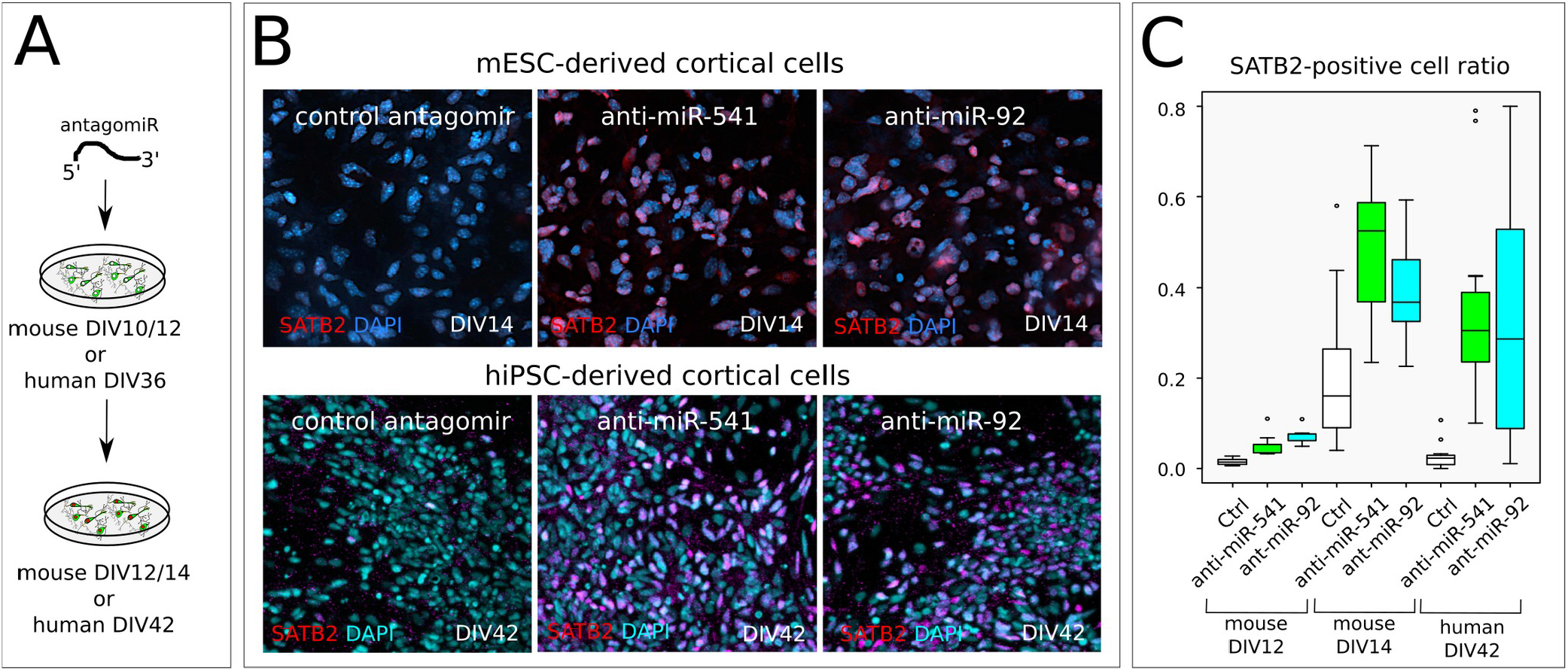
miR-92a-b and miR-541 function in mouse and human cortical cells. A, Outline of the *in vitro* assay of miR-541 inhibition by LNA-antisense oligonucelotide lipofection in corticalized mESCs (n= 2 independent experiments) or hiPSCs (n= 3 independent experiments). B, Immunocytodetection show SATB2-positive nuclei 2 days after mESC lipofection and 6 days after hiPSC lipofection, respectively. C, Box plots report SATB2-positive nuclei proportion. Ctr, scrambled sequence LNA lipofection. An anti-miR-92 LNA oligonucleotide was used to inhibit both miR-92a and miR-92b, which share the seed sequence.

### miR-541 targets are enriched in genes related to the development of supra-granular neurons

To infer the biological relevance of miR-92a/b and miR-541, we evaluated their degree of miRNA-mRNA target affinity using an *in silico* prediction approach (Enright et al., 2003). First, we analyzed the affinity of miRNA-*Satb2* 3’UTR interaction in relation to the average embryonic cortical miRNA expression of the mouse miRNAome. Among the annotated mouse miRNAs with significant affinity to *Satb2* 3’UTR (Supplementary file 12), miR-92a/b and miR-541 show high expression in cortical progenitors (miR-92a/b) or high *in silico* affinity to *Satb2* 3’UTR (miR-541) (Figure 7A), in line with their high miR-catch enrichment (Figure 5B). We then compared miR-92a/b and miR-541 targets with the targets of three recently described miRNAs of corticogenesis, namely let7, miR-9, miR-128 (Shu et al., 2019). To this aim, we selected a subset of 395 genes associated with an embryonic cortical marker signature (Galfrè et al., 2020). Among the 6 miRNAs analyzed, let-7 and miR-541 showed *in silico* affinity with more than half of the signature genes (Figure 7B, Supplementary File 13-14), suggesting for them a more relevant role in corticogenesis.

**Figure 7.**
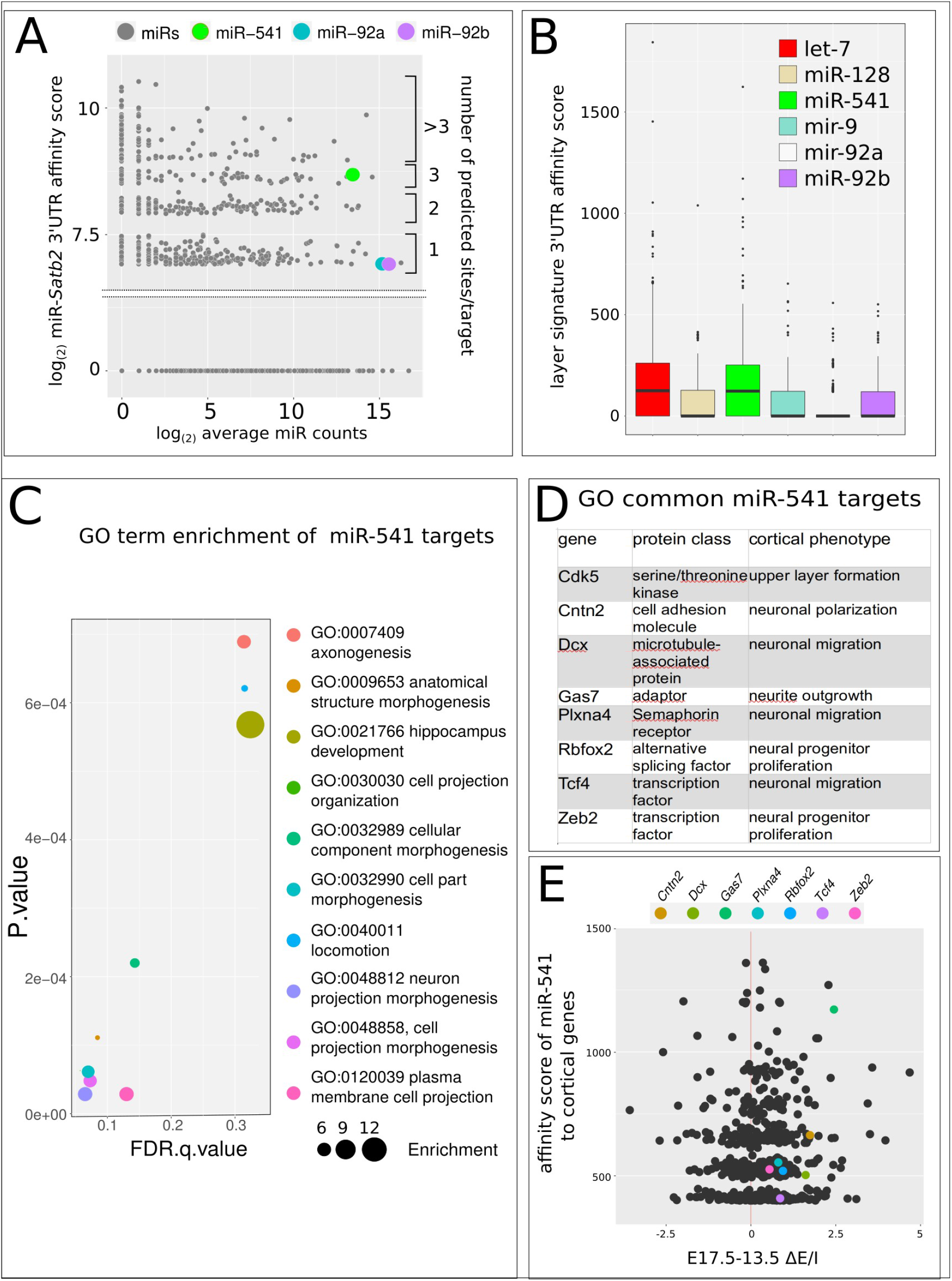
*In silico* analysis of miRNA/mRNA interactions. A, *In silico* comparison of the affinity of mouse miRNAome (grey dots), miR-92a-b and miR-541 (colored dots) to *Satb2* 3’UTR (Ensembl Mus musculus Satb2-201 cDNA 3’ UTR), in relation to the average miRNA expression levels during corticogenesis. B, *In silico* affinity of cortical miRNAs to the 3’ UTR of an embryonic layer gene signature (385 genes) (Galfrè et al., 2020). C, GO enrichment of the mir-541 gene targets with high *in silico* affinity to *Satb2* 3’UTR (cumulative score higher than 400, n=48) (Enright et al., 2003) with respect to the layer gene signature employed in B. D, list of the 8 genes common to all the GO terms shown in C. E, plot showing E/I read counts developmental increase (x-axis) with respect to miR-541/mRNA affinity score of genes of the embryonic cortical marker signature (Galfrè et al., 2020). Colored dots indicate genes listed in D. Names in labels indicate the 5 genes with the highest E/I read count ratio increase and mir-541/mRNA affinity score.

Interestingly, when analysing the mRNAs with the highest *in silico* affinity (total score higher than 400) for the 6 miRNAs, only the putative targets of miR-541 showed significant enrichment in GO terms (Figure 7C). It may be notable that terms related to neuronal projection development (axogenesis, neuron projection morphogenesis, cell projection morphogenesis, plasma membrane cell projection) (Figure 7C) are the most represented and that at least 8 out of the 11 putative target genes are related to cortical neuronal layering and migration, axon guidance, corpus callosum disturbances (Supplementary Figure 6). Interestingly, all these 8 genes are involved in basic processes controlling the generation of supra-granular layer cells such as polarization, proliferation and migration of late cortical progenitor cells (Caubit et al., 2016; Chen et al., 2016; Hatanaka et al., 2019; Li et al., 2019; Namba et al., 2014; Okamoto et al., 2013; Pramparo et al., 2010; Shinmyo et al., 2017; Ton and Kathryn Iovine, 2012; Zhang et al., 2016) (Figure 7D). Figure 7E compares the change of E/I read counts by EISA of 7 out of the 8 genes (not enough Cdk5r read counts were available for a significant analysis) to that of the genes of the embryonic cortical marker signature. The results indicate that all these 7 genes increase their E/I read count ratio between E 13.5 and E 17.5 and that there is a general correlation between E/I read count increase and mir-541/mRNA affinity score, supporting a relevant role of miR-541 in their post-transcriptional control during early corticogenesis. Altogether, these findings suggest a possible role of miR-541 as a hub in the post-transcriptional control of genes involved in the generation of supra-granular neurons.

## DISCUSSION

There is growing evidence that translational control exerted by RNA-binding proteins or miRNAs plays a crucial role in setting the appropriate time of production of key proteins involved in controlling the differentiation potential, the final layer of destination of cortical progenitors, as well as the differentiative program of the post-mitotic neurons (Kosik and Nowakowski, 2018; Nowakowski et al., 2018; Shu et al., 2019; Zahr et al., 2018). For example, cortical progenitors express Brn1 and Tle4 mRNAs, for both deep and superficial layer fates, respectively, but their translation into the corresponding proteins is initially repressed by a translational repression complex and subsequently released in due time (Zahr et al., 2018). Micro-RNAs are especially interesting in this respect and they have been indicated as heterochronic modulators of vertebrate development (Gulman et al., 2019; Robinton et al., 2019), also in the context of the vertebrate nervous system (Chiu et al., 2014; Nowakowski et al., 2018; Zahr et al., 2019).

SATB2 protein plays a central role in cortical neurogenesis, both in the early embryonic phase and at later postnatal stages. Inactivation of *Satb2* by conventional knockout leads to absence of corpus callosum and to a change in the projection abilities of upper layer projection neurons, that divert their trajectories to subcortical targets (Alcamo et al., 2008; Britanova et al., 2008). These data were integrated by conditional knockouts: when *Satb2* is inactivated early, callosal axons fail to form, and instead layer II-III neurons project subcortically or to the septum (Leone et al., 2015; McKenna et al., 2015; Srinivasan et al., 2012). On the other hand, when *Satb2* is inactivated at later stages, the corpus callosum appears intact, though there are consequences for plasticity and long-term memory storage (Jaitner et al., 2016). Furthermore, besides being involved in layer II-III callosal neuron specification, *Satb2* also plays a role in layer V subcortical projection neurons (Srinivasan et al., 2012). These data indicate that SATB2 acts in a multifaceted way that is both cell-context and time dependent, and indicate that precise control of its expression may be relevant for cortical development. Significantly, Paolino et al. (Paolino et al., 2020) have recently shown that accurate timing of SATB2 protein appearance in mouse is crucial for appropriate axonal projection of layer II-III neurons through the corpus callosum. In fact, while SATB2 protein is readily translated from its mRNA in the dunnart marsupial model (where layer II-III axons project through the anterior commissure and the CC is absent), in the mouse SATB2 protein appearance is delayed with respect to its mRNA expression (and axons go through the CC). Strikingly, anticipated SATB2 protein production in the mouse reroute layer II-III commissural axons toward the anterior commissure instead of the corpus callosum. This showed that a post-transcriptional control may be relevant in timing SATB2 protein appearance within the developing early placental neocortex (Paolino et al., 2020).

To get more insights into the early regulation of *Satb2* mRNA translation, we initially sought for evidence of differential stability of the mRNAs for *Satb2* and other key genes involved in mammalian corticogenesis, namely *Bcl11b*, *Cux1*, *Tbr1*, and *Fezf2*. By EISA, we found that among them only *Satb2* mRNA shows an increase, in both its stability and the rate of its transcription, that could be related to the delayed SATB2 protein appearance. Then, in a cell culture model of cortical differentiation, we have shown that the *Satb2* 3’UTR drives a significant translational inhibition of a GFP reporter at an early (DIV 12), but not at a late (DIV 18), stage of differentiation; and that *Satb2* 3’UTR is bound by the AGO/RISC complex in much a stronger way at an early (DIV 12) than at a late (DIV18) stage of *in vitro* differentiation, suggesting its regulation by miRNAs. We then have studied the expression of miRNAs in this same model and identified miR-92a, miR-92b and miR-541 as candidate to modulate SATB2 onset of translation, on the basis of their temporal dynamics of expression and, significantly, of miR-catch biochemical selection. We also showed that antagonizing these miRNAs anticipates the appearance of SATB2-positive cells in both mESCs and hiPSCs induced to cortical differentiation *in vitro*. While the antagonism of miR-92a/b might exert this effect by anticipating the translation of EOMES (TBR2), and then the differentiation of intermediate progenitor cell progeny expressing SATB2 (Bian et al., 2013; Nowakowski et al., 2013), miR-541 has no predicted binding sites on *Eomes* mRNA. Therefore, the effect of miR-541 on the onset of appearance of SATB2-positive neurons is directly due to its binding to *Satb2* 3’UTR. For this reason, and because of its peculiar taxon-specificity, we focused our attention onto miR-541 role in corticogenesis. In fact, unlike miR-92, let-7b, miR-128 and miR-9, and other miRNAs involved in cortical development (Chiu et al., 2014; Nowakowski et al., 2013, 2018; Shu et al., 2019; Zahr et al., 2018), which are evolutionarily conserved, mir-541 appeared recently during vertebrate evolution, being present only in Eutherian mammals (see below). miR-541 has not been deeply studied, except for a report showing its function in inhibiting neurite growth in PC2 cells (Zhang et al., 2011). miR-541 expression declines during corticogenesis in a temporal pattern opposite to that of SATB2 protein, and its presence in Eutherians, but not in Metatherians or any other vertebrates, suggests that it might be involved in the up mentioned heterochronic shift of SATB2 translation between dunnart and mouse (Paolino et al., 2020). Our demonstration that miR-541 can bind *Satb2* 3’UTR and inhibits its translation both *in vitro* and *in vivo* provides a molecular mechanism contributing to this heterochronic shift.

*Satb2* is present in all vertebrates (Sheehan-Rooney et al., 2010) and is expressed with other CITF genes in the dorsal telencephalon (pallium) of birds and reptiles, though with different patterns of mutual co-expression, as compared to the mammalian neocortex (Nomura et al., 2018; Tosches and Laurent, 2019). This suggests that the same CITFs have evolved different mechanisms of cell identity regulation in homologous telencephalic structures of different vertebrates (Cárdenas and Borrell, 2019; Nomura et al., 2018; Tosches and Laurent, 2019). Notably, in mammals *Satb2* has acquired a novel transcriptional control, due to the genomic insertion of a SINE sequence (AS021) carrying a new cortical-specific enhancer (Sasaki et al., 2008; Tashiro et al., 2011). In addition, in the early mammalian neocortex, SATB2 binds the *Bcl11b* promoter with high efficiency and prevents its expression, although at later stages LMO4 relieves this inhibition (Alcamo et al., 2008; Britanova et al., 2008; Harb et al., 2016). In contrast, in reptilian and avian pallial cells SATB2 and BCL11B are coexpressed, and SATB2 cannot silence *Bcl11b*, because of very inefficient binding of its cis-regulatory sequences (Nomura et al., 2018). By leading to differential expression of these two proteins in separate layers, this mechanism may increase cortical heterogeneity in the mammalian brain (Nomura et al., 2018). In addition, in higher primates, SATB2 appearance is delayed over an extended period, possibly crucial for cortical expansion, during which deep layer neurogenesis is balanced with the expansion of progenitor cells (Otani et al., 2016). Altogether, these observations indicate that tight temporal control and initial repression of SATB2 expression (Paolino et al., 2020) (present work) may hold a crucial role in pallial evolution. Together with other previously published data (Chiu et al., 2014; Nowakowski et al., 2013, 2018; Shu et al., 2019; Zahr et al., 2018), our results suggest that post-transcriptional mechanisms of regulation may be of great relevance for layering down the mammalian neocortex (Figure 8).

**Figure 8.**
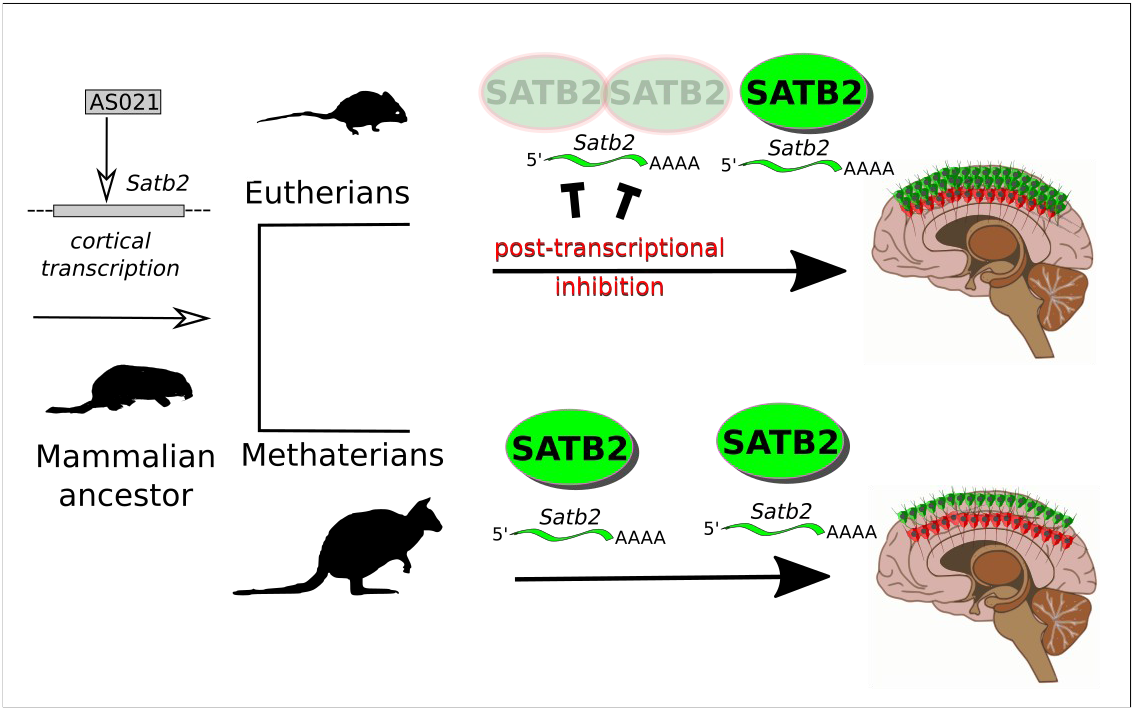
Post-transcriptional mechanisms and SATB2 heterochronic shift in mammalian brain evolution. The genomic insertion of a SINE sequence (AS021) carrying a new cortical-specific enhancer settled *Satb2* cortical expression in mammals (Sasaki et al., 2008; Tashiro et al., 2011). However, the control of SATB2 protein expression underwent a crucial change during mammalian evolution: a heterochronic shift occurred that may be involved in routing the axons of Eutherian supra-granular cell layers (green cells in the figure) to the corpus callosum (Paolino et al., 2020). miR-92a-b and miR-541, and possibly other post-transcriptional regulators, may have contributed to the shift by inhibiting SATB2 translation in the early corticogenesis phase.

miR-541 is encoded by *Mirg* (miRNA-containing gene), present only in Eutherian mammals inside the *Dlk1-Dio3* locus (Edwards et al., 2008; Glazov et al., 2008; Rocha et al., 2008; Winter, 2015). *Mirg* encodes about 40 miRNA genes (Glazov et al., 2008; Marty and Cavaillé, 2019). *Mirg* mRNA was detected in the developing early nervous system as well as in other organs, including the liver (Han et al., 2012). Constitutive *Mirg* deletion affects energy homeostasis causing neonatal lethality (Labialle et al., 2014), and behavioural disturbances (Lackinger et al., 2019; Marty et al., 2016). While the metabolic disorders have been related to alteration of liver gene expression program (Labialle et al., 2014), *Mirg* overall role, and of its individual miRNAs, in the early nervous system and in cortical layering has not been precisely defined, with few exceptions (Marty and Cavaillé, 2019; Winter, 2015). For some of these miRNAs a neurogenic function has been shown or proposed, but several seem involved in brain disorders (Gallego et al., 2016; Shi et al., 2015; Tsan et al., 2016; Winter, 2015). Furthermore, an overall GO analysis of the targets of these miRNAs pointed to embryonic and neural development and especially at axon guidance as key enriched terms; the possible involvement of *Mirg* in the regulation of key factors for formation of corpus callosum was suggested by in silico target analysis (Glazov et al., 2008). While we showed that miR-541 may play a role in timely control of SATB2 expression in upper cortical layer, it may also be notable that mRNAs for axon guidance molecules identified as targets of other miRNAs of *Mirg* (Glazov et al., 2008) are also in silico targets of miR-541; conversely, some of miR-541 most relevant targets (Supplementary Table 2) are also targets of other miRNAs of *Mirg*. The coordinate action of *Mirg* miRNAs may therefore be relevant in endowing the Eutherian brain with some of its characters.

## EXPERIMENTAL PROCEDURES

Mouse ES cell corticalization in vitro, cell transfection and analysis were performed as previously described (Terrigno et al., 2018a, 2018b). Human hiPS cells (ATCC-DYS0100 line, American Type Culture Collection) were neuralized according to Chambers at al. (Chambers et al., 2009).

Co-expression Table Analysis (COTAN) was performed on previously published datasets (Yuzwa et al., 2017) following the protocol of Galfrè et al. (Galfre et al., 2020). Exon-Intron split analysis (EISA) was performed as described (Gaidatzis et al., 2015; La Manno et al., 2018) on available datasets (Chui et al., 2020). RNA immunoprecipitation, Small RNA-seq and miR-catch were carried out as described (Marranci et al., 2019; Pandolfini et al., 2016), with minor modifications.

miRNA-mRNA *in silico* affinity was predicted as described (Enright et al., 2003), using score >120, energy < −18 kd as thresholds. 3’UTR sequences were obtained from Ensembl resources (Hunt et al., 2018), using Cran Biomart package. MiRNA sequences were obtained from miRBase database (v.22) (Kozomara et al., 2019). Detailed Material and Methods are described in Supplemental Information.

## AKNOWLEDGEMETS

We thank Giuseppe Lupo, Maria Antonietta Tosches, Richard Harland and Magdalena Gotz for critical review of the manuscript, Maria Antonietta Calvello and Edoardo Sozzi for technical support, Luciano Conti for IPSC neural differentiation methods advise, Austin Smith for ES cell lines. This work was supported by University and Research grant PRIN-2102 (F.C.), by the EU Commission, FP7 PAINCAGE Project, grant number 603191 and the H2020-ICT-2016 MADIA Project, grant number 732678.

## AUTHOR CONTRIBUTIONS

M.M., R.V. L.P. and F.C. designed the experiments; M.M. performed cell culture, molecular biology, imaging, and gene expression data computation; P.M. and I.A. planned and carried out IUE; S.G. and F.M. performed COTAN and EISA; M.P. and M.H.C. advised on computational methods of COTAN, EISA and small RNA-seq analysis; K.D. performed hIPSC cultures; M.T. and L.P. performed small RNA-seq; L.P. carried AGO RIP; M.R. and A.Marranci carried miR-CATCH; A.Mercatanti performed the computational analysis of captured miRNAS, R.V. and F.C. wrote the the paper.

