## Supplemental Information for "An Eutherian-Specific microRNA Controls the Translation of *Satb2* in a Model of Cortical Differentiation"

**Supplementary Material and Methods**

**Mouse ES cell-derived neural cell cultures**

Murine ES cell lines E14Tg2A (passages 25-38) and 46 C (transgenic *Sox1*-GFP ESC kindly provided by A. Smith, University of Cambridge, UK, passages 33–39) were used for *in vitro* corticalization. For expansion, ES cells were grown on gelatin-coated tissue culture dishes (pre-treated 10' with 0.1% gelatin in PBS) at a density of $4x10^4$ cells/$cm^2$. ES cell medium, changed daily, contained GMEM (G5154, Sigma-Aldrich), 10% Fetal Calf Serum (12133C, Sigma-Aldrich), 2mM Glutamine (25030, ThermoFisher Scientific), 1mM sodium Pyruvate (25030, ThermoFisher Scientific), 1mM non-essential amino acids (NEAA, 11140, Sigma Aldrich), 0.05mM β-mercaptoethanol (M3148, Sigma Aldrich), 100 U/ml Penicillin/Streptomycin (15140, ThermoFisher Scientific) and 1000 U/ml recombinant mouse LIF (PMC9484, ThermoFisher Scientific). Chemically defined minimal medium (CDMM) for neural induction consisted of DMEM/F12 (21331-046, ThermoFisher Scientific), 2mM Glutamine, 1mM sodium Pyruvate, 0.1Mm NEAA, 0.05mM β-mercaptoethanol, 100 U/ml Penicillin/Streptomycin supplemented with N-2 Supplement 100X (175020, ThermoFisher Scientific), and B-27 Supplement minus Vitamin A 50X (125870, ThermoFisher Scientific). ES neuralization was performed in three steps. In Step-I, dissociated ES cells were washed with DMEM/F12, seeded on gelatin-coated culture dishes ($6.5x10^4$ cells per $cm^2$) and cultured in CDMM plus 2.5µM 53AH Wnt inhibitor (C5324-10, Cellagen Technology) and 0.25µM BMP inhibitor (SML0559, Sigma Aldrich), for 3 days. In Step-II, ES cell were dissociated and seeded ($6.5x10^4$ cells per $cm^2$) on Poly-ornithine (P3655 Sigma-Aldrich; 20 µg/ml in sterile water, 24 hours coating at 37°C) and natural mouse Laminin (23017015, ThermoFischer Scientific; 2.5 µg/ml in PBS, 24 hours coating at 37°C). Cells were cultured for 4 additional days in CDMM plus Wnt/BMP inhibitors, with daily medium change. Serum employed for Trypsin inactivation was carefully removed by several washes in DMEM/F12. In Step-III, cells were dissociated and seeded ($1.25x10^5$ cells per $cm^2$) on Poly-ornithine and Laminin coated wells. Subsequently, isocortical culture were kept in CDMM Plus Wnt/BMP inhibitors for four additional days. On the eleventh day of differentiation, DMEM/F12 was replaced with Neurobasal and NEAA were removed from the CDMM to avoid glutamate-induced excitotoxicity. Medium was changed daily until the day of cell fixation.

To deplete cultures of post-mitotic cells, Cytarabin (AraC, 5 μM; AC449561000, ThermoFisher Scientific) was added to medium for two days before cell collection.

46 C *Sox1*::GFP cells were employed to identify progenitor cells at different times of corticalization *in vitro*. For fluorescence activated cell sorting of GFP-positive cells, $10^7$ cells were collected at each time of differentiation by trypsinization, washed, and resuspended in PBS, 2 % FBS, and 2 mM EDTA at a concentration of 10M cells per mL and kept on ice. The *Sox1*::GFP+ population was sorted on BD FACSJazz (BD Biosciences). Sorted cells ($10^5$ - $5x10^5$, depending on the culture stage), were immediately pelleted and total RNA extracted using miRNeasy Mini Kit (217004, Qiagen).

**hiPSC-derived neural cell cultures**

Neural cell cultures were differentiated from a commercial reprogrammed fibroblast line (ATCC-DYS0100 line, American Type Culture Collection). Cell neuralization was carried out essentially as described (Chambers et al., 2009), with minor modifications. Reprogrammed stem cells were seeded at $3x10^4$ cells/$cm^2$ cultured on 1:100 geltrex and maintained in Essential 8 medium for two days. After two days incubation, cultures were switched to neural differentiation media: DMEM/F12 1:1 (21331-046, ThermoFisher Scientific) containing 2mM Glutamine (25030, ThermoFisher Scientific), 1 mM Sodium Pyruvate (11360070, ThermoFisher Scientific), 100 U/ml Penicillin-streptomycin (15140, ThermoFisher Scientific), 1mM Non-essential amino acids (11140, Sigma Aldrich), 0.05mM β-mercaptoethanol (M3148, Sigma Aldrich), 10 μM 53AH (C5324-10, Cellagen Technology), 10 μM LDN193189 hydrochloride (SML0559, Sigma Aldrich), 1 μM RepSox (R0158, Sigma Aldrich), N-2

Supplement 100X (175020, ThermoFisher Scientific), and B-27 Supplement minus Vitamin A 50X (125870, ThermoFisher Scientific). After 10 days in neural differentiation medium, cells were displaced from substrate via incubation at 37ºC for 20 minutes in Accutase solution (A6964, Sigma Aldrich). Cells were harvested, diluted in 5 volumes of 1X PBS, centrifuged for 4 minutes, and replated at $10^5$ cells/cm$^2$ on poly-ornithine (P3655, Sigma Aldrich)/recombinant human laminin (AMS.892 021, Amsbio) in half volume of neural differentiation media + 5 μM Y-27632 (SM02, Cell Guidance Systems). Cells were maintained for 4 days in fresh neural differentiation media without ROCK inhibitor followed by an expansion of 7 days in neural differentiation media without TGFβ, WNT, and BMP inhibitors. After 11 days, cells were displaced again from substrate via incubation at 37ºC for 20 minutes in Accutase solution. Cells were harvested, diluted in 1X PBS at a volume 5 times that of Accutase, centrifuged for 4 minutes, and replated at 2.5x$10^5$ cells/cm$^2$ on poly-ornithine (P3655, Sigma Aldrich)/purified mouse laminin (CC095-M, Merck Millipore) in Eppendorf glass bottom dishes (H 0030 741 021, Eppendorf). Cells were maintained in neural differentiation media without inhibitors for 12 days and then switched to neuronal maintenance media based in Neurobasal (21103049, ThermoFisher Scientific) and containing 2mM Glutamine (25030, ThermoFisher Scientific), 1 mM Sodium Pyruvate (11360070, ThermoFisher Scientific), 100 U/ml Penicillin-streptomycin (15140, ThermoFisher Scientific), 0.05mM β-mercaptoethanol (M3148, Sigma Aldrich), Ascorbate, 0.5 mM (A92902, Sigma Aldrich), Recombinant human BDNF, 20 ng/ml (NBP2-52006, Novus Biologicals), and B-27 Supplement minus Vitamin A 50X (125870, ThermoFisher Scientific) until fixation at DIV 42. Cells were fixed with 2% PFA warmed to 37ºC for 15 minutes at room temperature.

**Cell transfection**

Plasmid transfections in mouse cortical cells were performed in 24-multiwell plate using 1 μg plasmid DNA diluted in 2.5 μL/well of Lipofectamine 2000 (12566014, ThermoFisher Scientific) in a final volume of 0.5 mL/well OPTI-MEM (31985062, ThermoFisher Scientific). Reporter activity plasmids were pEGFP-C1 (Clontech; control) and pEGFP-C1 fused to 3' UTR of *Satb2* between HindIII and XbaI sites.

LNA anti-miRNA (antagoMir) transfections in mouse cortical cells were performed using Lipofectamine 2000 according to the manufacturer's instructions. miRCURY LNA™ microRNA Inhibitors to miR-541-5p, miR92-3p and control antagomiR (MIMAT0003170, YI00199006 and MIMAT0000539, respectively) were resuspended in TE buffer (10 mM Tris pH 7.5, 1 mM EDTA) to a final concentration of 50 μM. Cells were transfected in 24-well plate using 25pmol of LNA diluted in 2.5 μL/well of Lipofectamine 2000 in a final volume of 0.5 mL/well OPTI-MEM.

After transfection, cells were incubated at 37°C and 5% CO2 for 4-6 hours and then the medium was replaced with complete Neurobasal medium (mouse cortical cells) or complete McCoy medium (HCT-116 cells).

##### *Satb2* 3'UTR cloning

The entire *Satb2*-3'UTR sequence (2802 bp) was obtained from Genome Reference Consortium Mouse Build 38 patch release 6 (GRCm38.p6) and amplified by PCR with Q5 High-Fidelity DNA Polymerase (M0491, NEB) with a forward and reverse primer carrying, correspondingly, a HindIII and XbaI restriction site at their 5'end (forward, CACAAAGCTTGTGAACTCCGCAGGCAGAGC; reverse, CACATCTAGAGCGTTTTATTTAACAACCAAAAAATTCTAACAGCC). The plasmid carrying the *Satb2*-3'UTR was constructed using mammalian expression vector pEGFP-C1 (Clontech) cut at HindIII position and XbaI positions inside the multiple cloning site and ligated with HindIII/XbaI restricted amplification product by T4 DNA ligase (M0202, NEB).

**Immunocytodetection (ICD) and imaging**

Cells prepared for immunocytodetection experiments were cultured on poly-ornithine/Laminin coated round glass coverslips. Cells were fixed using 2% paraformaldehyde for 12’, washed twice with PBS, permeabilized using 0.1% Triton X100 in PBS and blocked using 0.5% BSA in PBS for 1 hr at RT. Embryonic cortical sections were thawed and let dry at room temperature 1 hr, then briefly washed three times (5’ each) in PBS before antibody staining.

Cells/slices were pre-tretated 1 hr at room temperature with blocking solution: 1% BSA, 10% goat serum, 0.1% Triton X100 in PBS. Primary antibodies used for microscopy were SATB2 ab (1:1000; ab92446, Abcam), GFP ab (1:1000, ab13970, Abcam). Primary antibodies were incubated overnight at 4°C in PBS containing 1% BSA and 10% goat serum in PBS; cells/slices were then washed three times with PBS (10' each). Alexa Fluor 488 and Alexa Fluor 546 anti-mouse, anti-rabbit or anti-chicken IgG conjugates (1:500; A32723, A-11034, A-11039, A-11003, A-11010, A11040, Molecular Probes) were incubated 1 hour at RT in PBS containing 1% BSA and 10% goat serum, followed by three PBS washes (10' each). Nuclear staining was obtained with DAPI (D1306, ThermoFisher Scientific). Cells/slices were coverslipped with Aqua Poly-mount (18606-100, Polysciences).

Mouse neural cells were imaged using a Nikon Eclipse E600 epifluorescence microscope with a 20 X objective and a Photometrics Coolsnap CF camera. Five to ten optic fields from two or more biological replicates were acquired. In the experiments of EGFP pixel intensity quantification, all the pictures were acquired with the same parameters and the median of pixel intensity of the entire acquired field was analysed. For cell counting, double blind analysis was performed.

Human neural cells were imaged using a Leica SP2 confocal microscope with a 40X oil objective. Z-stacks were attained between 9-12 μm thick optical sections. Three biological replicates were attained per treatment group and subdivided into 5 technological replicate Z-stacks resulting in 15 total acquisitions. Stacks were flattened in ImageJ (RRID:SCR_002285) using the Z-stack projection function, set as a representation of standard deviation, and backgrounds were subtracted as a function of disabled smoothing and rolling ball radius of 20 px2. The resultant Hoechst+ and SATB2+ images were then subjected to an automated cell counter in ImageJ macros which analyzed separate channels at a 16-bit threshold set between 30-65355, and individual cells were counted using the "Analyze Particle" function set at circularity 35-150 px2 and circularity 0.33-0.99 to only include positive nuclei and minimize false positives.

**ScRNA-seq datasets**

ScRNA-seq datasets available in literature (Yuzwa et al., 2017) were used to analyse cortical gene expression at E11.5, E13.5, E15.5 and E17.5. Raw counts were obtained from GEO GSE107122 and used to plot red/cell values by the vioplot R package.

**COTAN**

Co-expression Table Analysis (COTAN) aims to estimate the UMI detection efficiency (UDE) of each cell, finds an approximation of the probability of zero read counts for a gene in a cell, and test the null hypothesis of independent expression for gene pairs, by counting zero/non-zero UMI counts in single cells (co-submitted paper). Briefly, mitochondrial genes and genes expressed in less than 0.3% of cells were eliminated. UDE for each cell and average expression for each gene were estimated as described (co-submitted paper) (*linear* method was used). PCA and hierarchical clustering (two clusters) were then carried out on UMI counts normalized dividing them by UDE. After removal of cell outliers resulting from PCA and hierarchical clustering, UDE and average expression were estimated again. Cells with very low UDE values were also removed. Together the two cleaning steps removed in all the datasets less than 3% of the cells (E11.5 dropped from 1,418 cells to 1,379 cells, E13.5 dropped from 1,137 to 1,119, E15.5 dropped from 2,955 to 2,921, E17.5 dropped from 880 to 863 cells).

Expected values for contingency table analysis were obtained as described (co-submitted paper) using cells UDE and genes average expression estimated with linear method, and genes dispersion estimated by fitting the observed number of cells with zero UMI count. COTAN then provided both an approximate p-value for the test of independence and a signed co-expression index (COEX), which measures the direction and intensity of the deviation from the independence hypothesis. The heatmaps in Figure 1D are colored by COEX value (blue for co-expression and red for disjoint expression).

For each gene, GDI was computed by normalizing P, the 0.001 quantile of the p-values of COTAN test for co-expression with all other genes. Our chosen normalization is ln(-ln(pval)). Genes with GDI > 2.2, which corresponds to $\ln(-\ln(10^{-4}))$, were generally non constitutive genes (co-submitted paper). Plots were generated with ggplot2 in R environment. The following R packages were employed: matrixStats, ggfortify, dplyr, rray, propagate, data.table, ggsci, gmodels, parallel, tibble, ggrepel.

**scRNA-seq bidimensional analysis**

UMI counts were divided by COTAN UDE for normalization. PCA was performed with normalized counts in R environment. Eigenvalues were plotted for selection by "elbow" point analysis (the number of components used were: 10 for E11.5, 10 for E13.5, 15 for E15.5 and 10 for E17.5). Selected components were employed as input for t-SNE function in sklearn.manifold python package (Loo et al., 2019), using the following parameters: perplexity 30, number of iterations 7000 and learning rate 700. Plots were obtained by ggplot2 R package.

**Exon-Intron split analysis (EISA)**

EISA on mouse cortex transcriptomes of cortical progenitor cells at E11.5, E13.5, E15.5 and E17.5 (Chui et al., 2020) was performed as previously described in (Gaidatzis et al., 2015; La Manno et al., 2018), with modifications. Mapping of datasets to mouse genome annotation GRCm38.98 was carried out as described in https://www.kallistobus.tools/velocity_index_tutorial.html (La Manno et al., 2018). Briefly, by using USCS table browser we obtained intron BED file, cDNA file and genome fasta files. A mouse GTF file was obtained from Ensembl. t2g utility and used to map transcripts to gene map (https://github.com/sbooeshaghi/tools/releases/tag/t2g_v0.24.0). Intron BED file was converted to fasta format by bedtools (v2.25; https://github.com/arq5x/bedtools2/releases). Association of intron and exon identifiers was performed modifying the fasta file headers as described in (https://www.kallistobus.tools/velocity_index_tutorial.html). An index was eventually produced by Salmon (version 1.1.0) (Patro et al., 2017) using the modified fasta files. Read pseudo-counts obtained by Salmon were normalized as reads per million (RPM). $Log_2$ (CPM) expression levels (exonic and intronic) were calculated and the exons/introns ratio was defined as the difference between $log_2$ exonic pseudo-counts and $log_2$ intronic pseudo-counts for each experimental condition.

**In Utero Electroporation (IUE)**

All animal procedures were approved by the internal Ethical Committee for Animal Experimentation (OPBA) of the Ospedale Policlinico San Martino and by the Italian Ministry of Health according to the Italian law D. lgs 26/2014 and the European Directive 2010/63/EU of the European Parliament. In all the experiments, the C57BL/6J strain from Jackson Laboratory was used.

In utero intraventricular electroporation was performed on E13 mouse embryos following laparotomy of deeply anesthetized pregnant females. Embryos were injected within the telencephalic ventricles with approximately 2µl (2µg) of pEGFP-C1 (Clontech; control) or pEGFP-C1 bearing normal *Satb2* 3' UTR, which were immediately electroporated at 35V with 4 pulses lasting 50 ms and spaced by 950 ms with a NEPA21 (NepaGene, Chiba, Japan) electroporator. Brains were dissected 7 days after electroporation and fixed overnight at 4°C in 4% paraformaldehyde in PBS. Brains were then cryoprotected overnight in 20% sucrose, embedded in Tissue Teck O.C.T. compound (4583, Sakura) and sectioned with a Leica CM3050 S cryostat at 12 µm thickness.

**RNA Immunoprecipitation**

Cross-linking Immunoprecipitation (CLIP) was carried out to enrich AGO-interacting RNA.

Cells were differentiated into cortical neurons until DIV12 or DIV18. Adherent cells were rinsed twice in PBS, cross-linked 150 mJ/cm$^2$ at 254 nm wave length, scraped, spun down 10 seconds at top speed and lysed on ice for 10' in 1 mL of fresh lysis buffer (Tris-HCl pH 8.0 25 mM, NaCl 150 mM, $MgCl_2$ 2 mM, 0.5% NP-40, DTT 5 mM) with protease inhibitors (1 tablet/10 mL lysis buffer of EDTA-free Complete Protease Inhibitor Cocktail Tablets, 11697498001, Sigma Aldrich) and RNasin (250 U/mL final, N2115, Promega). Cell lysate was centrifuged at 10000 rpm at 4ºC for 10' and the supernatant was kept at 4ºC for later procedure.

In the meantime, protein A Dynabeads (10001D, ThermoFisher Scientific) were rinsed 3 times with PBS/0.5% NP40 and incubated with 5 µg rabbit monoclonal Anti-argonaute-2 antibody EPR10411 (ab186733, Abcam), or anti-GFP antibody A-6455 (A-6455, ThermoFisher Scientific) in PBD/0.5% NP40 for 1 hour. After the initial binding, antibody-protein A beads were blocked with 0.5 mg/mL yeast RNA (10 µg/µL, 10109223001, Sigma Aldrich) and 1 mg/mL BSA (20 mg/mL, A3294-100G, Sigma Aldrich) for an additional 30'; beads were then washed twice in PBS/0.5% NP40 to remove the unbound IgGs and then twice in lysis buffer. The beads were resuspended in 100 µL of lysis buffer.

The lysate was subjected to preclearance by incubation with pre-blocked Protein A beads at 4ºC for 60' (100 µL of total lysate after pre-clearance, but before co-IP, was separated for total RNA – input – analysis). The remaining lysates proceeded to co-IP with anti-Ago-Protein A beads at 4ºC for 90'. After incubation, beads were washed three times with lysis buffer, twice with lysis buffer high-salt content (Tris-HCl pH 8.0 25 mM, NaCl 0.9 M, MgCl2 1mM, NP-40 1%, DTT 5Mm) and, again, once with lysis buffer.

After washes, beads were incubated with 100 µL of SDS 0.1% and Proteinase K (0.5 mg/mL, P8107S, NEB) for 15 minutes at 55ºC.

RNAs that co-immunoprecipitated with anti-AGO or anti-GFP antibodies were extracted adding 700 µL Qiazol (79306, Qiagen) and 140 µL chlorophorm according to manual and then purified using

Nucleospin RNA XS purification system (740902.50, Macherey-Nagel) and following manufacturer's instructions.

**Semiquantitative Real-Time PCR**

RNA quantity and quality was measured using NanodropTM Lite UV Visible Spectrophotometer (ThermoFisher Scientific) followed by reverse transcriptase protocol. For each sample, 100 ng of total RNA were reverse transcribed. Reverse Transcriptase Core kit (RT-RTCK-03, Eurogentec) was employed for cDNA synthesis. Primers for amplification were 5'CATGAGCCCTGGTCTTCTCT3' (*Satb2* forward) and 5'AACTGCTCTGGGAATGGGTG3' (*Satb2* reverse). Amplified cDNA was quantified using Sensi Fast SYBR Green (BIO-98050, Bioline) on Rotor-Gene 6000 (Corbett). Amplification take-off values were evaluated using the built-in Rotor-Gene 6000 "relative quantification analysis" function and relative expression was calculated with the 2-ΔCt method.

**Small RNA-Seq**

Total RNA was extracted with miRNeasy Mini Kit (217004, QIAGEN). Small-RNA libraries were prepared using TruSeq Small RNA Sample Preparation Kit (RS-200-0012/24/36, Illumina) following the manufacturer's instructions starting from 1μg of total RNA per sample. Libraries were multiplexed, loaded into a V3 flow cell and sequenced in a single-reads mode (50 bp) on a MiSeq sequencer (Illumina), obtaining ~4 million reads per samples. Raw sequences were demultiplexed to FASTQ format using CASAVA v.1.8 (Illumina). Quality control checks were performed with the FastQC algorithm. Adapters were trimmed from the primary reads using Cutadapt v1.2.1 (Martin, 2011). Remaining reads, with a length of between 17bp and 35bp, were clustered by unique hits and mapped to pre-miRNA sequences (miRBase release 21) (Kozomara and Griffiths-Jones, 2014) with the miRExpress tool v 2.1.3 5 (Wang et al., 2009). Read counts were CPM normalized for comparative analyses. PCA was carried out by PCA.GENES R package.

**miR-catch**

miR-catch analysis (version 2.0) (Marranci et al., 2019; Vencken et al., 2015) was performed essentially as described, with minor modifications. Three biological replicas for each time of *in vitro* differentiation were included in the study. Mouse cells (>$10^7$/sample) were harvested at DIV12 and DIV18 of the cortical differentiation protocol by trypsinisation, washed with PBS and fixed with 1% formaldehyde for 10 minutes at room temperature. The reaction was quenched with 1.25M glycine for 5 minutes at room temperature and cells were centrifuged at 200g for 5 minutes at 4°C. The pellet was resuspended in ice cold PBS (50 mL) and centrifuged at the same conditions as previously twice. Cells were then resuspended in 1ml Lysis Buffer (50mM Tris-HCl pH 7.0, 5mM EDTA, 1% SDS) plus supplements: 1mM Phenylmethanesulfonyl fluoride (PMSF, P7626, Sigma Aldrich), 1X Protease Inhibitor Cocktail (P8340, Sigma Aldrich) and 80U/ml RNAsin (N2115, Promega); all the components were added freshly before use. Cells were sonicated in ice-cold Lysis Buffer with a Soniprep 150 ultrasonic disintegrator (MSS150.CX3.1, MSE) for 12 rounds at 70% amplitude for 30 seconds pulses with 45 second cool down pauses in between. Sonicated lysates were pooled in order to have a minimum of 1ml to be hybridized with two probe pools, each containing 12 antisense oligonucleotides, odd or even, as indicated in Table 1.

| PROBE # | PROBE (5'-> 3') | PROBE POSITION * | PERCENT GC |
|---|---|---|---|
| **1** | aaagtccttggacccatcta | 24 | 45.0% |
| **2** | tctgagcttactcagtctat | 154 | 40.0% |
| **3** | cttccataagttggcaggaa | 273 | 45.0% |
| **4** | attgtaaagttctctgtccc | 408 | 40.0% |
| **5** | agtgactcactgtgaagtgg | 492 | 50.0% |
| **6** | attacccattaaaagctgcc | 627 | 40.0% |
| **7** | ctctggaggaattggtctta | 753 | 45.0% |
| **8** | ctcgatacagtgctggcatg | 835 | 55.0% |
| **9** | ggtccaacgtcaaaacgtca | 928 | 50.0% |
| **10** | gaaggaaagggtaacaccct | 1048 | 50.0% |
| **11** | tctaaccgggcagaaacttc | 1231 | 50.0% |
| **12** | tctggctaaagtgaagggga | 1336 | 50.0% |
| **13** | tcacttactttattgcctgg | 1441 | 40.0% |
| **14** | tggcattagttctgctttac | 1537 | 40.0% |
| **15** | ctggaaggtaatgctactgt | 1635 | 45.0% |
| **16** | tgctgagtggccatctcaag | 1724 | 55.0% |
| **17** | tgtattgcaacgtgtcttct | 1976 | 40.0% |

| | | | |
|---|---|---|---|
| **18** | gctcatgtcaagggtaactg | 2078 | 50.0% |
| **19** | ggagatcaggaagcagcaac | 2196 | 55.0% |
| **20** | agagtgacttcagcaacagc | 2245 | 50.0% |
| **21** | gatgccatcgatcgatgaac | 2310 | 50.0% |
| **22** | aaatgcccacagattcactt | 2436 | 40.0% |
| **23** | ctttgtcaagaggcactaca | 2557 | 45.0% |
| **24** | acagcctaacaatgcacata | 2739 | 40.0% |

**Supplemenray Table 1 miR-catch probes**

Dynabeads MyOne Streptavidin C1 (65001, TermoFisher Scientific) were washed (30 µl for each experiment) three times with 1ml unsupplemented Lysis Buffer and resuspended in 30 µl complete Lysis Buffer. The beads were added to 1ml lysate in a 1.5ml tube and kept on rotation in a 37°C hybridization oven for 30 minutes. Then, the lysates were cleared from beads twice using a magnetic stand and transferred to a 5ml round-bottom tube where 2ml of supplemented Hybridization Buffer (750mM NaCl, 1% SDS, 50mM Tris-HCl pH 7.0, 1mM EDTA and 15% formamide plus supplements: 1mM PMSF, 1X protease inhibitor cocktail and 80U/ml RNAsin that were added fresh before use) was added. At this point, a total amount of 100pmol probes (capture odd/even or scrambled control probes, 1 µl from a 100 µl pool previously mixed) were added to each lysate and put again in the 37°C hybridization oven for 4 hours in rotation. While the probes were incubating with the lysate, 200 µl of beads were washed three times with unsupplemented Lysis Buffer and resuspended in 200 µl supplemented Lysis Buffer. 100 µl of beads were added to the lysate plus probes sample and rotated in the hybridization oven for an additional 30 minutes at 37°C. After this, beads were pelleted using the magnetic support and resuspended in 1 ml of Wash Buffer (2X SSC Buffer, 0.5% SDS and 1mM PMSF added fresh) pre-warmed at 37°C. Five washes of 5 minutes each using hybridization oven in rotation at 37°C were performed with the Wash Buffer. At the last wash, the beads were spin down, the entire wash buffer was removed. Beads were then resuspended in 185 µl Proteinase K buffer (100mM NaCl, 10mM Tris-HCl pH 7.0, 1mM EDTA, 0.5% SDS) and then added with 15 µl 20 mg/ml Proteinase K and incubated at 45°C for 1 hour under constant and vigorous agitation followed by 10 minutes incubation at 95°C. Finally, 1 ml Qiazol was added directly to the beads, vortexed and incubated for 10 minutes at room temperature. The RNA extraction was performed using Nucleospin RNA XS purification system (740902.50, Macherey-Nagel).

The RNA eluted from the the ODD and EVEN samples were used to prepare cDNA libraries with the TruSeq Small RNA kit (RS-200-0012/24/36, Illumina), as per the manufacturer's suggestions. cDNA libraries were multiplexed, loaded into a V3 flow cell and sequenced in a single-reads mode (50 bp) on a MiSeq sequencer (Illumina), obtaining ~4 million reads per samples. Read counts were obtained as described in the miR-seq section method. To evaluate the enrichment of miRNA binding to *Satb2* 3'UTR, at each time of analysis (DIV12 or DIV18) *Satb2*-captured RNA from three biological replicas (DIV12: 3 EVEN and 2 ODD; DIV18: 2 EVEN and 2 ODD) and total RNA (DIV12 and DIV18, n=3) were compared. miRNA reads were normalized as CPM. Aiming to discover miRNAs with high biological relevance, those in the highest quartile of expression were considered for the analysis. The enrichment of miRNA binding to *Satb2* 3'UTR was evaluated as $\log_2$ captured/input fold change. The non-parametric NOISeqBIO statistical test of NOISeq R-package was applied with a probability >0.9 (Tarazona et al., 2015).

##### ***In situ* hybridization**

miRNA in situ hybridization (ISH) was performed using LNA-modified oligonucletides probes (Exiqon), according to the manufacturer protocol, with minor modifications, Cryosections were collected on slides (J1800AMNZT, Thermo Scientific) and postfixed 15' with 4% paraformaldehyde

(PFA) in PBS. Sections were treated with 10ng/µL proteinase K (15'), washed with 2 mg/mL glycine (2x 5'), PBS (2x5'), and postfixed 15' with 4% PFA. Sections were then pre-hybridized (50") in hybridization solution containing: with 50% formamide, 5X sodium saline citrate buffer (SSC) (pH 6), 1% sodium dodecyl sulfate (SDS), 50 g/mL heparin (9041-08-1, ThermoFisher Scientific) and 500 g/mL yeast RNA (10109223001, Sigma Aldrich). Hybridization with the digoxigenin-labeled probes was per-formed overnight at a temperature of approximately 21°C lower than the melting temperature of the probe. miRNA probes (miRNA Detection Probes, 339111, Exiqon) to mmu-miR-541-5p, and control probe with scrambled sequence, were employed. Washes were carried out in 50% formamide, 2XSSC at the hybridization temperature (1x 30') and 1X SSC (2x 15'). Sections were blocked 30' in MABT (1% BSA, A3294-100G, Sigma Aldrich; 150 mM NaCl; 0.1% Tween 20, pH7.5) containing 10% sheep serum (S2263, Sigma Aldrich) and incubated with alkaline phosphatase (AP)-labeled anti-digoxigenin antibody (1:2000; 11093274910, Sigma Aldrich)) in MABT and 1% BSA, overnight at 4°C. Sections were washed 5x5' in MABT and 3x5' in NMNT (100 mM NaCl, 100 mM TrisHCl pH 9.5, 50 mM MgCl, 0.1% Tween-20, 2 mM Tetramisole (L9756-5G, Sigma Aldrich: 500 mg/L). Sections were eventually with BM-Purple AP-substrate (L9756-5G, Sigma Aldrich) at RT 0.5'- 2 hours, then blocked by washes with PBS and counter-stained with anti-SATB2 antibody.

**MiRNA-mRNA interaction prediction and GO enrichment**

miRNA-mRNA *in silico* affinity was predicted as described (Enright et al., 2003), using score >120, energy < -18 kd as thresholds. 3'UTR sequences were obtained from Ensembl resources (Hunt et al., 2018), using Cran Biomart package. MiRNA sequences were obtained from miRBase database (v.22) (Kozomara et al., 2019). Enriched GO terms were obtained using two unranked lists of genes (target versus background) as described (Eden et al., 2009) Analysis results were visualized using Cran ggplot2 packages.

### Supplementary Figures

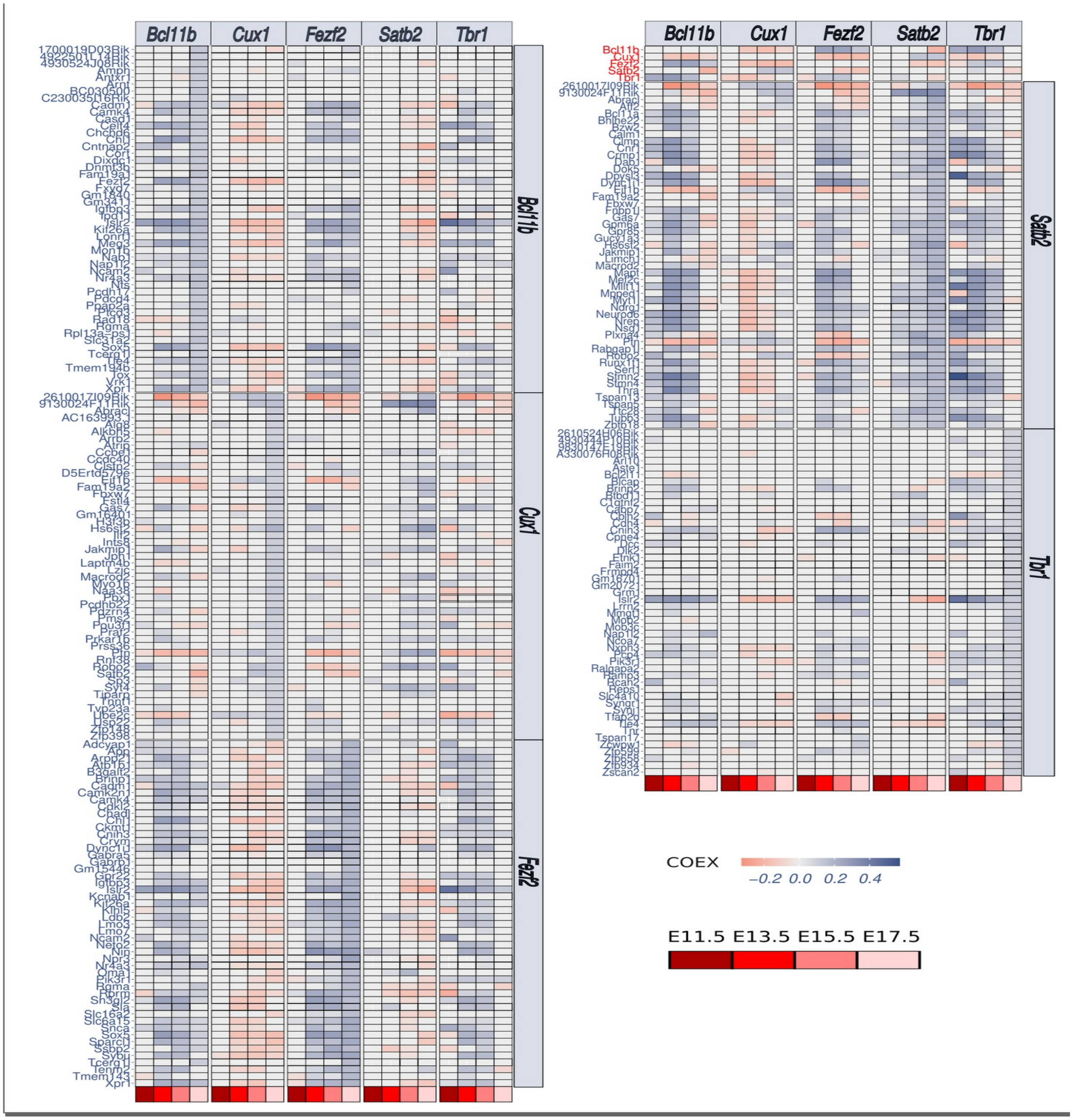

**Supplementary Figure 1 COEX values of genes related to CITFs.** Heatmap shows details of main Fig.1 D.

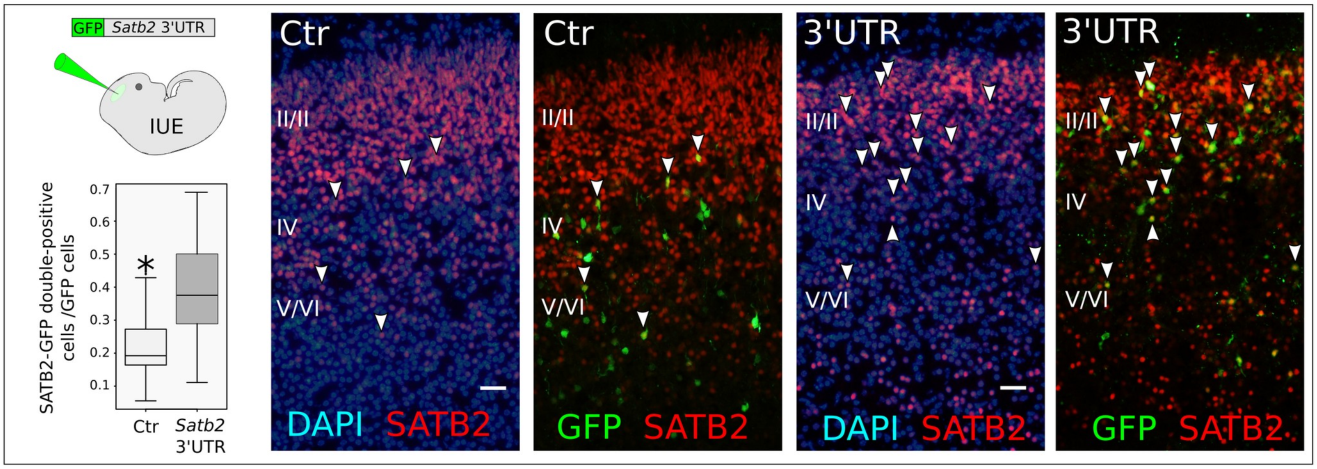


**Supplementary Figure 2 Expression of a GFP reporter bearing Satb2 3' UTR**. IUE at E13.5 of a reporter as in the scheme. Box plot indicates the ratio of GFP-SATB2 double-positive cells out of all GFP-positive cells. Pictures show examples of electroporated cells at E19. Arrowheads point to GFP-SATB2 double positive cells. Roman numbers indicate layers. Scale bar, 50 μm. Data from n= 3 animals for control IUE and n=3 animals for 3' UTR IUE.

DIV12

| miRNA | C_mean | WT_mean | theta | prob | log2FC |
|---|---|---|---|---|---|
| mmu-miR-9-5p | 19162.9172355122 | 144170.598307786 | -1.65275053829413 | 0.999999999823203 | -2.91138786671142 |
| mmu-miR-127-3p | 36455.6956630299 | 65206.8940538959 | -0.662647575691103 | 0.972458927466191 | -0.838880270755658 |
| mmu-miR-370-3p | 1616.61203109597 | 54209.89031013 | -1.96077260376643 | 0.997598637007709 | -5.06751069303321 |
| mmu-miR-99a-5p | 260.023479173067 | 35788.1934247691 | -2.06625406429111 | 0.992162416975841 | -7.10469800939344 |
| mmu-miR-381-3p | 592.995740973867 | 30257.0572889963 | -2.10249945965394 | 0.995916192827015 | -5.67310612888758 |
| mmu-miR-541-5p | 55663.5734970947 | 26348.4377597595 | 0.955364524165008 | 0.989390928562448 | 1.07901610521601 |
| mmu-miR-99b-5p | 14047.1160699048 | 23288.4527849742 | -0.346114557554273 | 0.886168381259914 | -0.729340824713096 |
| mmu-miR-181a-5p | 57902.9806393449 | 19053.6002671297 | 1.07886460035826 | 0.99268019030525 | 1.60357398746084 |
| mmu-miR-30d-5p | 10858.4354699307 | 16595.5608643057 | -0.332586869951845 | 0.872497119472575 | -0.611981139213528 |
| mmu-miR-30a-5p | 11465.7782840027 | 16314.6094647321 | -0.309665565282424 | 0.84533092920688 | -0.508830166207279 |
| mmu-miR-26a-5p | 8080.04839674217 | 13071.1040460943 | -0.379213909766239 | 0.9133144904286 | -0.693945163628539 |
| mmu-miR-434-5p | 1226.33527990417 | 11435.835599681 | -1.00208421868079 | 0.993672541134473 | -3.22113641450054 |
| mmu-miR-103-3p | 4912.91438769687 | 10615.3577129697 | -0.455942227783417 | 0.949403756556448 | -1.1115019834852 |
| mmu-miR-129-5p | 446.764842385 | 9124.69188071557 | -1.19566495598532 | 0.998458807173419 | -4.35218827902783 |
| mmu-miR-125b-5p | 5457.1597293008 | 8841.74567769397 | -0.24003424693085 | 0.719783023416249 | -0.696180965934634 |
| mmu-miR-134-5p | 812.3420888572 | 8347.820338382 | -1.02080454350702 | 0.99375374810244 | -3.36124025120868 |
| mmu-miR-92a-3p | 72846.6496936747 | 7828.73543763937 | 3.27925713381272 | 0.999999999999999 | 3.21801142714565 |
| mmu-let-7i-5p | 6470.7580425965 | 7289.5696372048 | -0.070890751564195 | 0.176537587071276 | -0.171898910655777 |
| mmu-miR-300-3p | 446.764842385 | 6489.32498700907 | -0.980626439560605 | 0.994135806916537 | -3.86048085408463 |
| mmu-miR-298-5p | 35087.1062515417 | 6171.71742632153 | 1.21136245561836 | 0.999007915901907 | 2.50719705449791 |
| mmu-miR-21a-5p | 4843.26600746753 | 6079.50857535097 | -0.100335885859878 | 0.285730207258889 | -0.327974468592859 |
| mmu-let-7c-5p | 8030.54752770413 | 5968.4259329594 | 0.152130849168717 | 0.22261363926849 | 0.428147858646983 |
| mmu-miR-92b-3p | 175527.646975037 | 5815.67129573947 | 2.54781245904834 | 1 | 4.91560874412339 |
| mmu-miR-744-5p | 8628.68764168573 | 5528.34974372723 | 0.268611426971269 | 0.608316200632184 | 0.642292264248633 |
| mmu-miR-25-3p | 8974.54196831753 | 5416.95923491583 | 0.315559916930315 | 0.674654441586653 | 0.728355076398944 |
| mmu-miR-181d-5p | 5298.31641360817 | 5327.18097169433 | -0.003102904675557 | 0 | -0.007838288549519 |
| mmu-miR-128-3p | 333.1389284675 | 5241.02212045253 | -1.07533965459178 | 0.995496886404853 | -3.97565234506373 |
| mmu-miR-433-3p | 4176.23655940263 | 5069.13661418287 | -0.118175905839089 | 0.351503284766943 | -0.279536610885456 |
| mmu-miR-770-3p | 439.026029099967 | 5028.08712869457 | -0.762183828637822 | 0.992145970691877 | -3.51763126724626 |
| mmu-miR-296-3p | 519.880291679433 | 4971.7803774281 | -0.802351009850374 | 0.994324616003707 | -3.25751119868855 |
| mmu-miR-151-3p | 8847.0509247046 | 4918.4304228307 | 0.346462111202041 | 0.714529258714192 | 0.846998633549695 |
| mmu-miR-106b-3p | 406.254377761933 | 4843.1659567017 | -0.939457118176282 | 0.995230881781305 | -3.57549517482481 |
| mmu-miR-488-3p | 73.4487826277667 | 4748.82763940873 | -1.22361874930652 | 0.999272229153218 | -6.0146890046085 |
| mmu-miR-434-3p | 14905.2351555125 | 4652.59796020197 | 0.738844391349579 | 0.968639873765927 | 1.67971070490009 |
| mmu-miR-148a-3p | 3304.81081761313 | 4639.05339147273 | -0.14421167724899 | 0.442189074908108 | -0.489262762771824 |
| mmu-miR-423-3p | 892.065797964667 | 4320.62481813267 | -0.623078724743014 | 0.967760385505086 | -2.27601792874398 |
| mmu-miR-181b-5p | 16010.0846705562 | 4278.63178980287 | 0.777227835565773 | 0.970356624305905 | 1.90375950361084 |
| mmu-miR-30e-5p | 4142.33435459263 | 4213.2009293873 | -0.008410208525823 | 0.007428130091215 | -0.024472712791209 |
| mmu-miR-409-3p | 3536.93737072867 | 3843.76374242837 | -0.045561548975601 | 0.092417624027704 | -0.12001899017873 |
| mmu-miR-125a-5p | 5800.58740378367 | 3812.45608242823 | 0.230566812705134 | 0.514903475633794 | 0.605478285427849 |
| mmu-miR-328-3p | 1072.1989013636 | 3804.74382499767 | -0.504458873583853 | 0.959899043498835 | -1.82722675778881 |
| mmu-miR-382-5p | 333.1389284675 | 3751.68235387233 | -0.913976581483621 | 0.995163628631504 | -3.49334183040838 |
| mmu-miR-218-5p | 73.4487826277667 | 3729.9681630927 | -1.09067870162138 | 0.995820282137518 | -5.66628092715414 |
| mmu-miR-149-5p | 1039.5939166922 | 3615.0254227171 | -0.446220294375599 | 0.946408693052749 | -1.79798569613988 |
| mmu-miR-100-5p | 673.201393484033 | 3612.4213136175 | -0.661510623544632 | 0.97224251330126 | -2.4238560948535 |
| mmu-miR-342-3p | 771.831624234133 | 3607.00906887417 | -0.564295675770599 | 0.966429287314849 | -2.22444498870297 |
| mmu-miR-140-3p | 479.369827056367 | 3500.92585223507 | -0.71844133716191 | 0.985284465319257 | -2.86852549707125 |
| mmu-miR-5099 | 32512.7841638476 | 3463.9335316699 | 1.53022995949927 | 0.999975168750474 | 3.23052395144994 |
| mmu-miR-340-5p | 666.1111902683 | 3256.61684540593 | -0.586482970492011 | 0.967100491870377 | -2.28953906697247 |
| mmu-miR-93-5p | 1137.4088707063 | 3244.7868806858 | -0.375713564046559 | 0.910837850786843 | -1.51237276327027 |
| mmu-miR-125b-1-3p | 2152.1366160122 | 3241.4649434234 | -0.156003526355483 | 0.480937664281697 | -0.590876308200271 |
| mmu-miR-30c-5p | 2143.41585932447 | 3218.47031497127 | -0.185666475846565 | 0.572960263427075 | -0.586463377773568 |
| mmu-miR-191-5p | 6459.92366757233 | 3001.33186395257 | 0.413171107626217 | 0.808937021310173 | 1.10591426793357 |
| mmu-miR-9-3p | 73.4487826277667 | 2981.15904666987 | -1.00286677353922 | 0.993665994334611 | -5.34299095670397 |
| mmu-let-7g-5p | 925.9680027747 | 2862.62041339687 | -0.434515830686556 | 0.942289599533236 | -1.62830213356896 |
| mmu-miR-320-3p | 26477.0394522772 | 2764.5836607757 | 1.35790011194303 | 0.994316681605955 | 3.25960767951089 |
| mmu-miR-323-3p | 219.679681216633 | 2714.4307456592 | -0.720816744159017 | 0.985786277432472 | -3.62717633411346 |
| mmu-miR-532-5p | 73.4487826277667 | 2692.24377483693 | -0.988795778122872 | 0.993905248128206 | -5.19592665835248 |
| mmu-miR-148b-3p | 146.5642319222 | 2588.57512253867 | -0.803110933819202 | 0.994339047283877 | -4.14255321608084 |
| mmu-miR-501-3p | 731.4878262777 | 2218.4145965911 | -0.32411428556095 | 0.86308014805803 | -1.60062325432953 |
| mmu-miR-760-3p | 1876.46884360237 | 2151.24503431923 | -0.05174185365066 | 0.111443482544014 | -0.197151525829117 |
| mmu-miR-210-3p | 146.5642319222 | 2019.64157030313 | -0.638238322609558 | 0.968859650981708 | -3.7844943069144 |
| mmu-miR-99b-3p | 146.5642319222 | 1892.52582293687 | -0.681919748285854 | 0.976656131650518 | -3.69070801457886 |
| mmu-miR-666-5p | 479.369827056367 | 1778.9615421243 | -0.452774279271559 | 0.948468046550294 | -1.89182431303167 |
| mmu-miR-431-5p | 1591.91252637637 | 1764.81450390223 | -0.03923200851678 | 0.074193493517858 | -0.148755488943228 |
| mmu-miR-3099-3p | 9640.5512989871 | 1708.45288847093 | 0.802859587141407 | 0.969800708187724 | 2.49642518473231 |
| mmu-miR-423-5p | 18196.9398895762 | 1609.7072801599 | 1.21557113841961 | 0.99915252028513 | 3.49882559018772 |
| mmu-miR-27b-3p | 6722.59893907447 | 1544.38665230687 | 0.585685691299704 | 0.90772915460315 | 2.12198509230751 |
| mmu-miR-17-5p | 226.936551099033 | 1504.05519491717 | -0.511389413845414 | 0.960968153263166 | -2.728496613407 |
| mmu-miR-30a-3p | 519.880291679433 | 1473.68408845583 | -0.345131588454824 | 0.885228469683108 | -1.50317591936386 |
| mmu-miR-101a-3p | 333.1389284675 | 1460.2760320085 | -0.495330881481169 | 0.958363019483719 | -2.13204525058758 |

DIV18

| miRNA | C_mean | WT_mean | theta | prob | log2FC |
|---|---|---|---|---|---|
| mmu-miR-9-5p | 17181.7040725625 | 118689.159298949 | -1.40941628509275 | 0.999999999493706 | -2.78824313525393 |
| mmu-miR-370-3p | 1074.12099569315 | 90529.5973389199 | -2.19109388008768 | 1 | -6.3971611157717 |
| mmu-miR-127-3p | 31134.1623914373 | 63582.902145437 | -0.765630020450287 | 0.978436642233718 | -1.03014040098699 |
| mmu-miR-129-5p | 1100.65722238685 | 61090.5640236974 | -2.31805793451028 | 1 | -5.79451241474891 |
| mmu-miR-99a-5p | 502.7646438116 | 52681.4200062846 | -2.55733488659366 | 1 | -6.71126722524581 |
| mmu-let-7i-5p | 15622.8349861939 | 51434.410716591 | -1.08701383347854 | 0.998209174317141 | -1.71907760132551 |
| mmu-let-7c-5p | 25803.9041265353 | 26139.8915568557 | -0.013863004759952 | 0.002005663069475 | -0.018663794335631 |
| mmu-miR-125b-5p | 6862.65093135172 | 22761.6112576957 | -0.920182415686784 | 0.995337402803095 | -1.72976480831467 |
| mmu-miR-30d-5p | 11975.5751644936 | 20257.2631020122 | -0.433215136661351 | 0.860056914360728 | -0.758344321538567 |
| mmu-miR-128-3p | 1259.16947342487 | 18069.1153887999 | -1.74670956003907 | 1 | -3.84298150215707 |
| mmu-miR-26a-5p | 10724.204448226 | 14785.2934010962 | -0.261534201047356 | 0.636263607995832 | -0.463292244250308 |
| mmu-miR-181a-5p | 72448.9567520552 | 14650.2159429583 | 1.35451417193874 | 0.990227114202529 | 2.30604298545381 |
| mmu-let-7g-5p | 1721.3642384427 | 14434.2153276876 | -1.22399619708568 | 0.999880874954577 | -3.06786837477338 |
| mmu-miR-30a-5p | 8486.88455676925 | 12082.9004873404 | -0.262561623176434 | 0.638698055903864 | -0.50965985526101 |
| mmu-miR-99b-5p | 8493.8155749798 | 11073.1219877182 | -0.203780767412904 | 0.475643626049409 | -0.382577347938065 |
| mmu-miR-103-3p | 5115.71721301512 | 10873.4110699992 | -0.426267937980226 | 0.855502515231092 | -1.08779617340814 |
| mmu-miR-770-3p | 265.9557365439 | 10761.1278640024 | -1.288300540259 | 0.999994870229456 | -5.33849932757877 |
| mmu-miR-433-3p | 2379.60717697462 | 10636.059102193 | -0.74546640481347 | 0.976547236278142 | -2.16016835932688 |
| mmu-miR-541-5p | 15800.9805349703 | 10482.2679091608 | 0.312223310899234 | 0.53177353354766 | 0.59206320066719 |
| mmu-miR-328-3p | 1478.51633244413 | 9316.8116696166 | -0.870778378747773 | 0.991276746147707 | -2.65568615061988 |
| mmu-miR-381-3p | 683.465617120875 | 9157.8273828221 | -1.08998761030961 | 0.998219632561889 | -3.74406470411935 |
| mmu-miR-7b-5p | 51.66757283545 | 8720.72511912283 | -1.54381241800847 | 1 | -7.3990451758275 |
| mmu-miR-100-5p | 1115.67159225973 | 8505.5358453962 | -0.993529780760716 | 0.998075656999002 | -2.93048970827859 |
| mmu-miR-434-5p | 530.72486358905 | 7182.92414875033 | -1.16191954944849 | 0.999214639818594 | -3.75853523748777 |
| mmu-let-7f-5p | 14348.8357022891 | 6814.77968295047 | 0.526277539158943 | 0.829063343170036 | 1.07419475646043 |
| mmu-miR-298-5p | 23812.0868519338 | 6312.62909159913 | 0.991015102804321 | 0.983297837752125 | 1.9153811706587 |
| mmu-miR-744-5p | 4188.30463793505 | 6098.3471501473 | -0.194704482473657 | 0.446403560506031 | -0.542051897451986 |
| mmu-miR-125a-5p | 4344.60434352677 | 6087.20474854643 | -0.19110465577115 | 0.434567576925784 | -0.486555093972103 |
| mmu-let-7b-5p | 2221.7139753338 | 5676.8951156567 | -0.478321064536765 | 0.889193629037971 | -1.35342899178927 |
| mmu-miR-760-3p | 1318.06217274495 | 5607.72938518413 | -0.594737757320744 | 0.953668039375622 | -2.08899830661507 |
| mmu-let-7a-5p | 17948.0097221012 | 5547.47669249993 | 1.09906820527611 | 0.994080466539033 | 1.69392026448552 |
| mmu-miR-92a-3p | 58375.4189275634 | 5430.2634757987 | 2.38438363433744 | 0.999999999999999 | 3.42626689402225 |
| mmu-miR-5099 | 18981.5322906205 | 4463.01135530477 | 1.03653330533842 | 0.987859795721606 | 2.08850707672853 |
| mmu-miR-92b-3p | 122546.705568659 | 4446.3518936604 | 3.34275642969298 | 0.999999999999999 | 4.78456575962324 |
| mmu-miR-379-5p | 51.66757283545 | 4238.34625372577 | -1.21865717277889 | 0.999853551445089 | -6.35809853144892 |
| mmu-miR-344-3p | 71.19652974505 | 4095.8420468751 | -1.20801832063254 | 0.999784432652282 | -5.84620934652477 |
| mmu-miR-382-5p | 124.312677053825 | 4004.29281241803 | -1.05679968360506 | 0.998244588882987 | -5.00950214361905 |
| mmu-miR-148a-3p | 4350.14707305922 | 3928.5302723957 | 0.048969731741832 | 0 | 0.14707449996477 |
| mmu-miR-423-3p | 860.0054705105 | 3840.36092900543 | -0.610236138681935 | 0.958909033882208 | -2.15882416429717 |
| mmu-miR-320-3p | 31740.2507348892 | 3705.0908501836 | 1.3429081593941 | 0.988755852479388 | 3.09873270142603 |
| mmu-miR-181d-5p | 4417.28069494687 | 3705.0785452893 | 0.096253152229803 | 0.058398664579533 | 0.253654383905084 |
| mmu-miR-181b-5p | 23386.9781441297 | 3620.4327057586 | 1.40420428569115 | 0.994501228462111 | 2.69147142129305 |
| mmu-miR-151-3p | 7905.14835635342 | 3426.9061323684 | 0.510978474390821 | 0.815876958585972 | 1.2058858629608 |
| mmu-miR-125b-1-3p | 2844.86047969973 | 3392.31322064513 | -0.082917258292857 | 0.105756436693641 | -0.253911483370619 |
| mmu-miR-434-3p | 18574.0606252739 | 3377.54186099163 | 1.31186367082689 | 0.984885708313994 | 2.4592436935341 |
| mmu-miR-3099-3p | 2649.4716130871 | 3100.02054087663 | -0.067250104165379 | 0.074401134621145 | -0.226573104722662 |
| mmu-let-7e-5p | 7117.60849636658 | 2977.89839678987 | 0.448459753384973 | 0.748242101223911 | 1.25709804837005 |
| mmu-miR-191-5p | 11827.6371248153 | 2937.62815030617 | 0.931126485842134 | 0.976991813187976 | 2.00943819305982 |
| mmu-miR-488-3p | 323.855997773925 | 2679.71376037183 | -0.774604509318561 | 0.979444967235366 | -3.04865453623103 |
| mmu-miR-300-3p | 451.47207097615 | 2666.43049617727 | -0.651575721355583 | 0.968590174383363 | -2.56220107548154 |
| mmu-let-7d-3p | 355.494613699775 | 2578.42775002203 | -0.628995631262896 | 0.964039812304788 | -2.85859201439416 |
| mmu-miR-106b-3p | 210.8910147618 | 2559.26502555833 | -0.808825470679418 | 0.983852602306892 | -3.60116001990617 |
| mmu-miR-21a-5p | 9021.42412887957 | 2518.77653958723 | 0.624614746229869 | 0.899775774636547 | 1.84063206143343 |
| mmu-miR-501-3p | 424.790207497275 | 2435.9376169228 | -0.533197716402333 | 0.923802851308921 | -2.51965477323776 |
| mmu-miR-30c-5p | 1378.24567508328 | 2435.25798787053 | -0.262327876164345 | 0.638145032384666 | -0.821241543247408 |
| mmu-miR-149-5p | 1492.47262306995 | 2433.25582078447 | -0.206950584043085 | 0.485630342059465 | -0.705183538360776 |
| mmu-miR-140-3p | 559.52794006705 | 2414.2570823159 | -0.533291728496342 | 0.923857940659811 | -2.10929723055287 |
| mmu-miR-30e-5p | 3811.65874442592 | 2364.7077383897 | 0.23725878030318 | 0.37526132557473 | 0.688757073988394 |
| mmu-miR-25-3p | 6105.1191386863 | 2328.95440701003 | 0.583704592084322 | 0.874909455363663 | 1.39033705226665 |
| mmu-miR-1981-5p | 191.3620578522 | 2309.39190471623 | -0.837997136462049 | 0.987619354836767 | -3.59313630563509 |
| mmu-miR-340-5p | 1832.67565135335 | 2263.39498604127 | -0.075816423060721 | 0.090953589786716 | -0.304536892957877 |
| mmu-miR-296-3p | 366.592307741475 | 2157.7800423674 | -0.62170110337163 | 0.962198018416143 | -2.55729938927821 |
| mmu-miR-134-5p | 226.772822724725 | 2143.24225539647 | -0.765478013943329 | 0.978420461672608 | -3.24047527227128 |
| mmu-miR-210-3p | 260.360013123925 | 2066.59407682043 | -0.70047153808044 | 0.97371027028103 | -2.98867524332212 |
| mmu-miR-532-5p | 35.785764872525 | 2049.69756326317 | -1.11370240357847 | 0.998418594772897 | -5.83988142719007 |
| mmu-let-7j | 2563.36668799115 | 2046.0208361894 | 0.100257515290508 | 0.065000217514516 | 0.325219032788286 |
| mmu-miR-342-3p | 362.94515879485 | 1828.28616539117 | -0.561335900199308 | 0.939160365902915 | -2.33266842248917 |
| mmu-miR-1839-5p | 983.177652338 | 1772.3922765258 | -0.210276557514178 | 0.495975740489693 | -0.850173916495047 |
| mmu-miR-323-3p | 311.6525859594 | 1765.6516180611 | -0.592231567391548 | 0.952732508627902 | -2.50219012313663 |
| mmu-miR-132-3p | 212.839589235125 | 1751.27323511313 | -0.59218170643163 | 0.952713610976892 | -3.0405657623655 |
| mmu-miR-124-3p | 276.241821086875 | 1666.96150183727 | -0.590833476354193 | 0.952199016703113 | -2.5932171307181 |
| mmu-miR-5121 | 717.05280752005 | 1645.77468261743 | -0.303836769014468 | 0.724149611035033 | -1.1986155592727 |
| mmu-miR-423-5p | 10720.0526282725 | 1645.6515893467 | 0.956478494364258 | 0.98043733151293 | 2.70358115679693 |
| mmu-miR-409-3p | 2299.7985557704 | 1548.7769886992 | 0.183459909335122 | 0.242069600089871 | 0.57037807526769 |
| mmu-miR-101a-3p | 564.413996852025 | 1528.56862949497 | -0.332889092949792 | 0.769760996242508 | -1.43735565687549 |
| mmu-miR-383-5p | 550.25382049865 | 1472.45977303843 | -0.373355074336348 | 0.815413846724438 | -1.42005905866972 |
| mmu-miR-148b-3p | 621.177293107475 | 1468.0622452298 | -0.348103485792116 | 0.789120213962276 | -1.24083614049895 |
| mmu-let-7d-5p | 2024.89717103298 | 1316.16517929217 | 0.165561396862178 | 0.198699333825979 | 0.621508087183256 |
| mmu-miR-93-5p | 1420.71801407788 | 1156.4516243103 | 0.079114273363537 | 0.032714844053662 | 0.296915317385665 |
| mmu-miR-329-5p | 35.785764872525 | 1147.42676925877 | -0.917593458914685 | 0.995171408226546 | -5.0028724554364 |
| mmu-miR-378a-3p | 1762.29892117258 | 1136.38131172803 | 0.168087100644849 | 0.204699186152204 | 0.633011643867364 |
| mmu-miR-652-3p | 364.17587656765 | 1103.72540497573 | -0.320868870983783 | 0.752319352874833 | -1.5996740249285 |
| mmu-miR-27b-3p | 9434.60480410963 | 1090.9465812036 | 0.980277415988635 | 0.982489750224922 | 3.1123816265705 |
| mmu-miR-337-5p | 18.08038243625 | 1057.1726799863 | -1.0612688432746 | 0.998228337549293 | -5.86964204350139 |
| mmu-miR-146b-5p | 1603.0304748744 | 1055.59750317313 | 0.153305653038486 | 0.170302238445163 | 0.602742009180353 |
| mmu-miR-690 | 599.82476172455 | 1000.73434971643 | -0.150027803821334 | 0.295292535561366 | -0.738446068578776 |
| mmu-miR-431-3p | 481.3578080907 | 980.779763731333 | -0.280211473630804 | 0.678205994970915 | -1.02681952000058 |
| mmu-miR-344f-3p | 18.08038243625 | 965.7768370087 | -1.04557584200134 | 0.998287222308045 | -5.7391927635529 |
| mmu-miR-135a-1-3p | 662.11308573795 | 956.180964963467 | -0.12605205694946 | 0.218157692911046 | -0.530206042257537 |
| mmu-miR-666-5p | 1942.4433394881 | 947.3204597402 | 0.252628228308036 | 0.411165969172406 | 1.03594806773227 |
| mmu-miR-708-3p | 493.663162769 | 907.938802496367 | -0.183046963839049 | 0.407667166501914 | -0.879068064348151 |
| mmu-miR-543-3p | 51.66757283545 | 857.243611415233 | -0.718777681095845 | 0.97477338829334 | -4.05237422993026 |
| mmu-miR-99b-3p | 628.2076200278 | 815.938311717533 | -0.074082089167849 | 0.08748515229657 | -0.377218639971437 |
| mmu-miR-1198-5p | 297.594352469775 | 811.3833511982 | -0.304994838768983 | 0.726196074144934 | -1.44703655185722 |
| mmu-miR-127-5p | 51.66757283545 | 782.706430399567 | -0.70007353339207 | 0.973685378546244 | -3.92114027989931 |
| mmu-miR-672-5p | 1468.75240893962 | 778.086029497433 | 0.256796235758521 | 0.420599310167882 | 0.91658963639103 |
| mmu-miR-129-2-3p | 140.06948501675 | 771.610871935633 | -0.557484361180435 | 0.937212760143372 | -2.46173078081615 |

**Supplementary Figure 3 MirCATCH analysis.** Analysis of RNAseq data from miRCATCH performed with NOIseqbio at the two indicated times (DIV12 and DIV18). Reported miRNAs are in the quartile of the highest expression. log2FC indicates the fold change between captured and input miRNAs. Means refer to average CPM of captured (C) and input (WT) miRNAs. Prob is the probability of differential expression calculated by NOIseqbio (Tarazona et al., 2015). Theta indicates differential expression statistics.

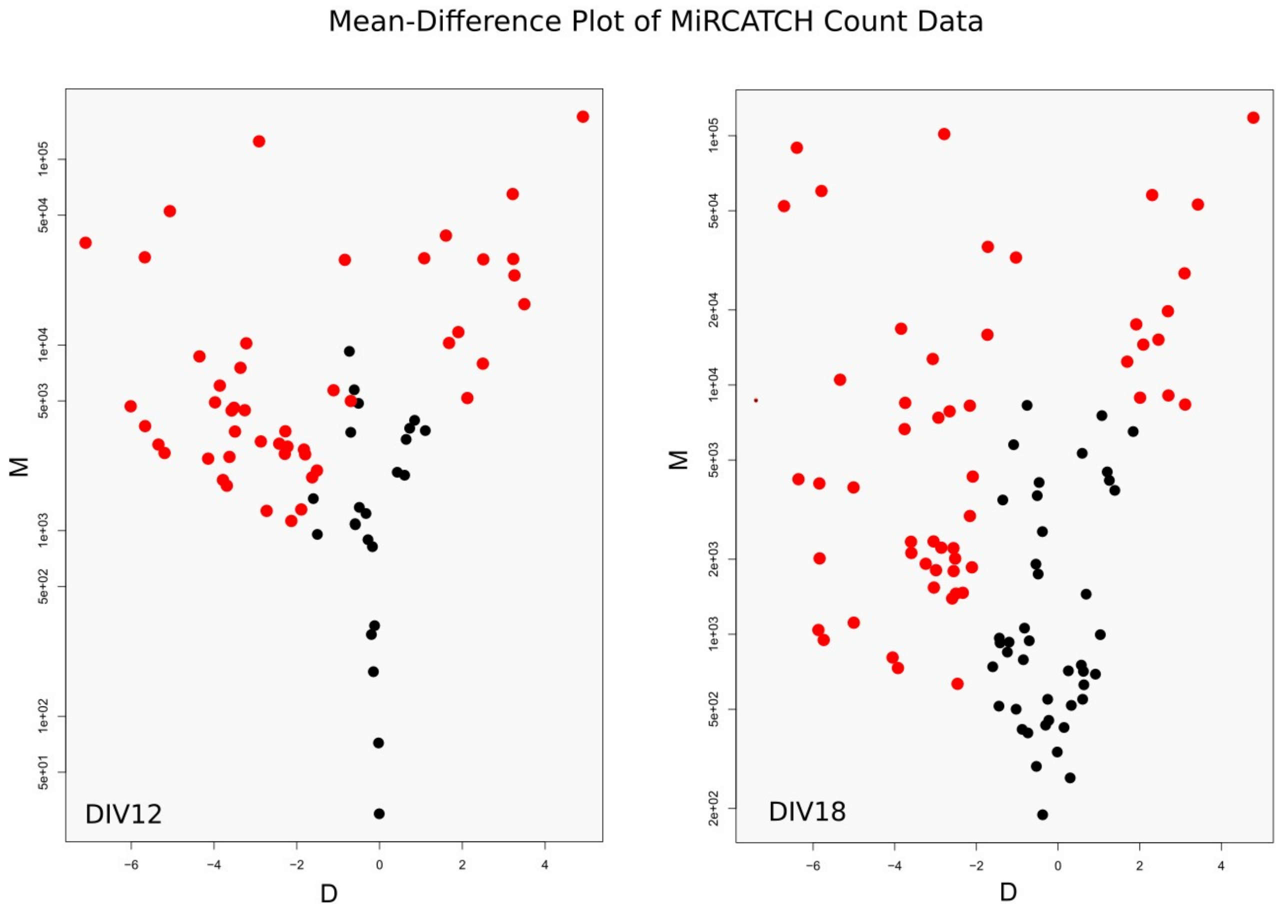


**Supplementary Figure 4 miR-CATCH M-D plot.** The Mean (M) - Difference (D) plot reported in figure shows the results of miRNA capturing by miRCATCH as obtained by analysis with the NOIseqbio R package. Plots show the distribution of miRNAs (as in Supplementary Figure 1) enriched (positive $\log_2$ fold change, D) or depleted (negative $\log_2$ fold change) after miRCATCH capturing, with respect to average expression (M, mean CPM), at the indicated time of differentiation (DIV12, DIV18). In red, miRNAs with significant fold change (probability > 0.9).

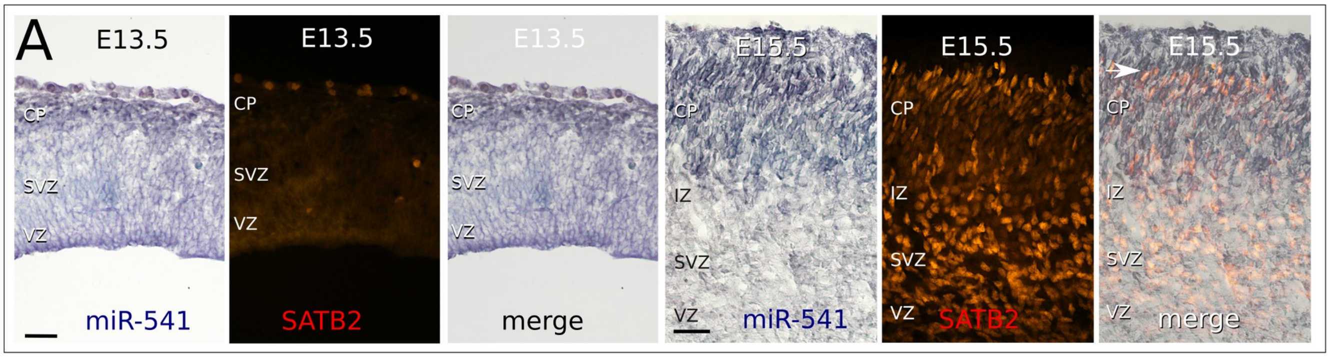


**Supplementary Figure 5 mir-541 expression**. *In situ* hybridization shows miR-541 distribution (BM-purple staining) compared to SATB2 immunodetection (red fluorescence). CP, cortical plate; IZ, intermediate zone; SVZ, subventricular zone; VZ, ventricular zone. Arrow indicates newly migrated SATB2-positive cells. Scale bar, 50 μm.

| Gene name | Biochemical role | Biological process | Phenotype/ disease | References |
|---|---|---|---|---|
| Gas7 | Adaptor protein | Neurite outgrowth, growth arrest | AD, schizophrenia | Gotoh et al., 2013; Ju et al., 1998; You and Chao, 2010; Zhang et al., 2016; |
| Rbfox2 | RNA binding protein | Neuronal mRNA splicing | | Underwood et al. 2005; Yeo et al., 2007; Jacko et al., 2018 |
| PlexinA4 | Semaphorin receptor | Cortical neuronal migration, axon guidance | | Hatanaka et al., 2019; Suto et al., 2005 |
| Cntn2 | GPI-linked member of the immunoglobulin superfamily, cell adhesion molecule | Neurite outgrowth, axonal adhesion | Smaller cortex, reduced corticothalamic axons, callosal and commissural defects | Kastriti et al., 2019 |
| Dcx | Brain-specific microtubule associated protein | Cytoskeletal regulator involved in migration, cortical layering, neurite extension | Lissencephaly, subcortical band heterotopia, epilepsy, cognitive disability | des Portes et al., 1998; Gleeson et al., 1998; Shahsavani et al., 2017 |
| Zeb2 | Transcription factor | Regulation of gene expression | Mowat-Wilson syndrome; defects in axonal growth; hippocampal and callosal defects | Takagi et al., 2015; Srivatsa et al., 2015 |
| Cdk5 | Serine/threonine kinase | phosphorylation of various cytoskeletal proteins | Corticogenesis, neuronal migration, axonal projections, dendrite branching; callosal dysgenesis | Ohshima et al., 1996; Gilmore et al., 1998; Ohshima et al., 2007; Hirota et al., 2007 |
| TCF4 | Transcription factor | Regulation of gene expression | Pitt-Hopkins syndrome; schizophrenia; autism; callosal dysgenesis | Amiel et al., 2007; Hasi et al., 2011; Forrest et al., 2018; Schoof et al., 2020 |

**Supplementary Table 2 miR541 target genes enriched in GO analysis**

Table references:


- Amiel J et al., 2007, Am J Hum Genet 80, 988-993. doi: 10.1086/515582
- des Portes et al., 1998, Cell 92, 51–61. doi: 10.1016/s0092-8674(00)80898-3
- Forrest et al., 2018, Schizophrenia Bulletin 44, 1100–1110. doi:10.1093/schbul/sbx164
- Gilmore et al., 1998, J Neurosci 18, 6370-6377. doi: 10.1523/JNEUROSCI.18-16-06370.1998
- Gleeson et al., 1998, Cell 92, 63–72. doi: 10.1016/s0092-8674(00)80899-5.
- Gotoh et al., 2013, J Biol Chem 288, 34699–34706. doi: 10.1074/jbc.M113.513119.

- Hasi et al., 2011, Hum Genet 130, 777–787. DOI 10.1007/s00439-011-1020-y
- Hatanaka et al., 2019, iScience 21, 359–374. doi: 10.1016/j.isci.2019.10.034
- Hirota et al., 2007, J Neurosci 27, 12829 –12838. doi:10.1523/JNEUROSCI.1014-07.2007
- Jacko et al., 2018, Neuron 97, 853–868. doi.org/10.1016/j.neuron.2018.01.020
- Ju et al., 1998, PNAS, 95, 11423–11428. doi: 10.1073/pnas.95.19.11423
- Kastriti et al., 2019, Front Cell Neurosci 13: 454. doi: 10.3389/fncel.2019.00454
- Ohshima et al., 1996, Proc Natl Acad Sci USA 93, 11173-11178. doi: 10.1073/pnas.93.20.11173
- Ohshima et al., 2007, Development 134, 2273-2282. doi:10.1242/dev.02854
- Schoof et al., 2020, Eur J Neurosci 00, 1–17. DOI: 10.1111/ejn.14674
- Shahsavani et al., 2017, Molecular Psychiatry 00, 1–11. doi:10.1038/mp.2017.175
- Srivatsa et al., 2015, Neuron 85, 998-1012. doi: 10.1016/j.neuron.2015.01.018.
- Suto et al., 2005, J Neurosci 25, 3628 –3637. DOI:10.1523/JNEUROSCI.4480-04.2005
- Takagi et al., 2015, Human Molecular Genetics 24, 6390–6402. doi: 10.1093/hmg/ddv350
- Underwood et al., 2005, Mol Cell Biol 25, 10005–10016. doi:10.1128/MCB.25.22.10005–10016.2005
- Yeo et al., 2007, PLoS Comput Biol. 3, 1951-1967. DOI: 10.1371/journal.pcbi.0030196
- You and Chao, 2010, J Biol Chem, 285, 11652–11666. doi: 10.1074/jbc.M109.051094.
- Zhang et al., 2016, Molecular Brain 9, 54. doi: 10.1186/s13041-016-0238-y.
